# Deep learning-based decoding of axonal ultrastructure in gene-edited mice using electron microscopy imaging

**DOI:** 10.64898/2026.05.26.727755

**Authors:** Nianqi Deng, Guillaume Miao, Hooman Bagheri, Alan Peterson, Anmar Khadra

## Abstract

Myelin forms an insulating sheath around axons enabling both rapid and energy-efficient conduction of action potentials and myelin abnormalities or loss can lead to severe motor, sensory, and cognitive impairment. While electron microscopy can resolve multiple axonal components that are affected myelin, their large-scale quantitative analysis is both difficult and time consuming. To overcome such limitations, we developed a machine learning framework that automatically recognizes and quantifies multiple features of axons and myelin including axonal mitochondrial density and periaxonal area. Applying that framework to fibers in the spinal cord of variably hypomyelinated mice, we show here that reduction in the thickness and length of myelin sheaths results in correlating changes in mitochondrial density and periaxonal area. The machine learning framework introduced here should contribute to future insight into the axon, myelin, and mitochondrial relationships that change during neurological plasticity and myelin disease progression.

## Introduction

Myelin is a multilayered membrane sheath surrounding axons that enables rapid propagation of electrical signals along myelinated nerve fibers. By insulating axons and enabling saltatory conduction, myelin accelerates signal transmission while reducing metabolic cost, thereby supporting efficient long-range communication across the central and peripheral nervous systems (1–4). Beyond conduction properties, myelin also contributes to neural plasticity, with activity-dependent oligodendrocyte dynamics enabling adaptive remodeling of myelin sheaths throughout life (5–9).

Disruption of myelin integrity has widespread consequences, impairing signal synchrony and axonal stability, and is implicated in diverse neurological disorders including multiple sclerosis, traumatic injury, schizophrenia, and Alzheimer’s disease (10–16). Despite its importance, how myelin thickness and composition vary across axon calibers, genetic backgrounds, and developmental stages remains incompletely understood, particularly in the context of mutations that alter myelin elaboration and stability (17, 18).

Myelin basic protein (MBP), encoded by the *Mbp* gene, comprises a major fraction of myelin proteins and is a central determinant of myelin compaction by mediating adhesion between the cytoplasmic surfaces of apposed myelin membranes (19–21). Loss of MBP prevents formation of compact myelin and disrupts formation of the major dense line. *Shiverer* mice lack MBP and, despite successful initiation of membrane wrapping they lack compact myelin (7, 22–26). This structural abnormality is accompanied by severe conduction deficits and neurological dysfunction (25).

Recently, models bearing different deletions in the *Mbp* transcriptional enhancers were derived. As different lines accumulate *Mbp* mRNA to different levels rather than complete deletion, these provide a stepwise framework for studying *Mbp* function (27). Progressive reduction in *Mbp* mRNA expression leads to increased g-ratio (defined as the ratio of axon diameter to fiber diameter) and accumulation of structural abnormalities while the intrinsic periodicity of myelin lamellae is unaffected (28). Such models enable systematic investigation of how incremental changes in myelin architecture affect multiple fiber properties including the density, ultrastructure and organization of axonal components.

Previous studies of fiber ultrastructure in *Mbp*-deficient models have revealed characteristic abnormalities including loss of myelin compaction, expansion or disruption of the periaxonal space (PAS), and increased axonal mitochondrial content within axons (7, 29–31). Such alterations in structural and metabolic organization demonstrate the importance of the myelin-axon relationship. However, the quantiation of such ultrastructural features remains quite challenging.

Spinal cord white matter exhibits substantial morphological heterogeneity, with densely packed axons of different calibers interspersed with vascular elements and processes of multiple glial cell types (32). Although axons, periaxonal space, myelin, and organelles are identifiable by their ultra-structural features in cross-sections, dense packing and local variability in fiber morphology make it difficult to consistently delineate individual fibers and adjacent periaxonal space or organelles. Manual annotation is therefore labor-intensive, limited in scalability, and prone to observer bias, particularly when quantifying large datasets or subtle phenotypic differences. Deep learning approaches have substantially advanced automated segmentation of axonal structures. The AxonDeepSeg algorithm provides robust two-class segmentation of axons and myelin across multiple imaging modalities (33, 34), while AimSeg and DeepACSON extended segmentation to additional compartments and three-dimensional reconstructions (35, 36). More recent methods such as Mysh improved per-fiber segmentation accuracy through cascaded architectures (37). Despite these advances, existing models are typically limited to a small number of classes, lack unified multi-compartment predictions, and often exhibit reduced performance when applied to genetically perturbed tissue, reflecting a persistent domain gap (38, 39). Moreover, they do not explicitly capture the coupled yet distinct relationships among axonal geometry, myelin structure, and organelle distribution (40, 41).

In this study, we introduce MicroStructFormer, a transformer-based framework that enables multi-compartment segmentation of spinal cord ultrastructure by combining fine-grained ultrastructural predictions with axon-myelin segmentation. MicroStructFormer identifies mitochondria-like organelles, PAS, and myelin inner tongue abnormalities, while AxonDeepSeg provides axon and myelin masks. These complementary predictions were integrated into a unified segmentation map spanning the major ultrastructural components.

Here we performed large-scale, quantitative analysis of axonal and myelin ultrastructural features in spinal cord white matter across the different *Mbp* enhancer-edited lines at multiple ages. This allowed us to systematically investigate how graded *Mbp* levels reshape myelin structure, axonal caliber distributions, and mitochondrial organization, providing a means to characterize how molecular perturbations of myelin are associated with large-scale alterations in axonal ultra-structure.

## Methods

### White matter ultrastructural dataset

Experimental data were obtained from a previously published study (28). Briefly, multiple mouse lines carrying distinct *Mbp* enhancer deletions or sequence alterations were generated via CRISPR editing in mouse zygotes. Quantiation of *Mbp* mRNA in the spinal cords of young mice revealed graded levels of *Mbp* mRNA as detailed in (27). The lines bearing the deleted or mutated enhancers were identified as: wild type (WT), M3KO, M3(225)KO, M5KO, M3M5KO, M1EM3M5KO, and M3KOKI. Knockout is abbreviated hereafter as KO, and knockin is abbreviated hereafter as KI. M3KO, M3(225)KO, and M5KO are collectively referred to as single KO; M3M5KO as double KO; and M1EM3M5KO as triple KO. Electron micrographs of cervical spinal cord cross sections of such mice provided the biological foundation for all downstream computational modeling and morphometric analyses.

### Overall strategy and rationale

An experimental workflow was used to generate EM images from different regions of the spinal cord (Fig. 1A). Tiled electron micrographs were obtained across three spinal cord white matter domains: corticospinal tracts (CST), gracile and cuneate tracts (GC), and ventromedial (VM), at three developmental ages: P30, P90, and P400 (Fig. 1B).

**Fig. 1.**
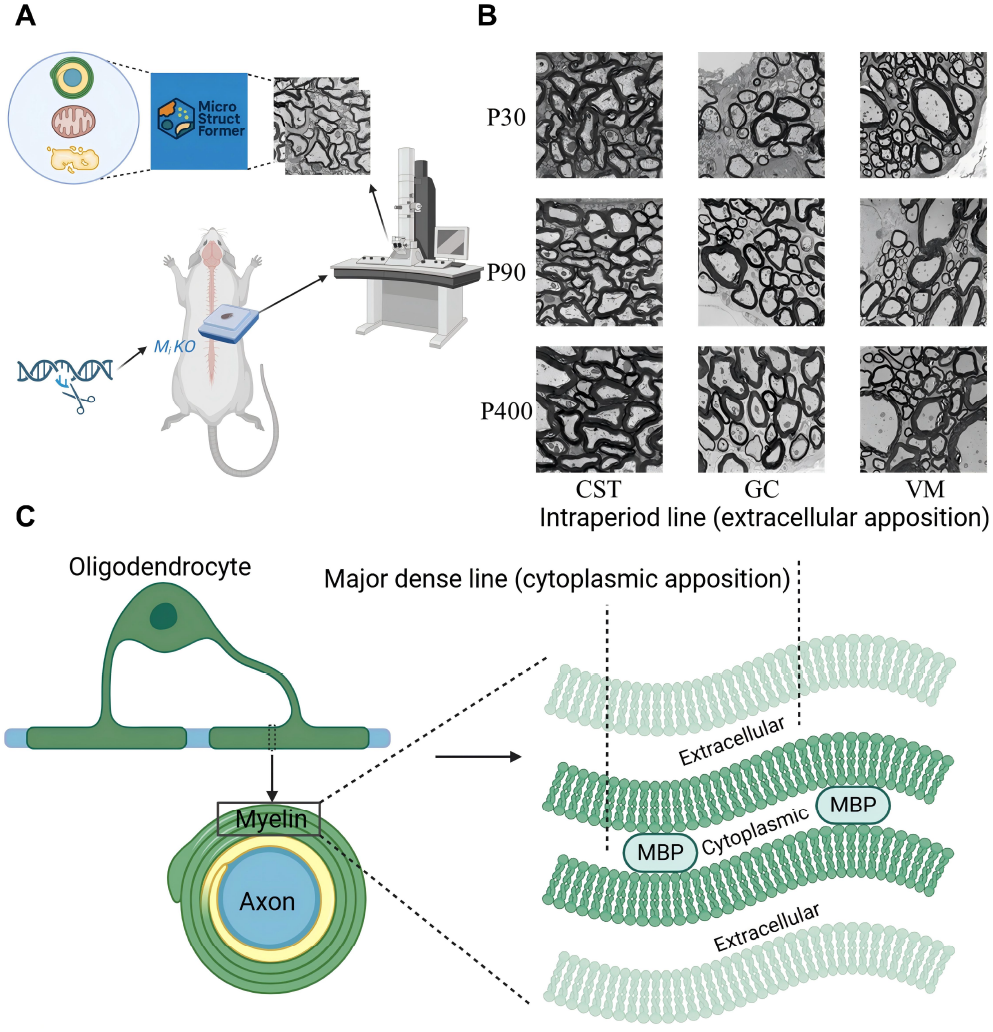
Schematic diagram highlighting the type of data set used in the analysis. Panels A and C were created in BioRender. Khadra, A. (2026) https://BioRender.com/iauaqmo. **(A)** Overview of the experimental workflow used to collect EM images from WT and enhancer-edited *Mbp* mouse lines. **(B)** Raw imaging data arranged by age and region. Only WT images are displayed for clarity, but all enhancer-edited *Mbp* mouse lines were included in the quantitative analyses. **(C)** Illustration of compact myelin ultrastructure and molecular organization. It is generated by oligodendrocytes that repeatedly wrap their membrane around axons, producing alternating cytoplasmic and extracellular leaflets. During compaction, MBP brings cytoplasmic leaflets into tight apposition to form the major dense line, whereas adhesion between extracellular leaflets produces the intraperiod line. Through iterative wrapping and MBP-dependent cytoplasmic compaction, these lamellae mature into the compact myelin that ensheathes axons.

These images served as the basis for developing a deep learning segmentation pipeline (MicroStructFormer combined with AxonDeepSeg), enabling automated multi-compartment segmentation and subsequent large-scale morphometric analysis of axons, compact myelin (Fig. 1C), mitochondria, and periaxonal space in relation to hypomyelination severity.

### Model initialization and domain-specific pre-training

MicroStructFormer was initialized from the Hugging Face model facebook/mask2former-swin-large-ade-semantic, which had been pre-trained on large-scale natural image datasets, providing strong generic feature extraction and class discrimination capabilities. To adapt it to electron micrographs, we conducted a domain-specific pre-training stage on publicly available EM datasets encompassing mitochondria, periaxonal space, and other organelles (35, 42). This two-stage pre-training was essential for bridging the substantial domain gap between natural and EM images and for reducing the burden of manual annotation.

### Dataset annotation and fine-tuning

Following pre-training, we fine-tuned MicroStructFormer on our own multi-genotype, multi-age, and multi-region spinal cord EM dataset. Initial annotations were manually created based on prior morphological literature (7, 29, 30, 43–46) and used as ground-truth segmentations (Fig. 2A). Because the dataset included gene-edited mouse lines exhibiting abnormal morphologies, we implemented an iterative, expert-guided annotation process, in which preliminary model predictions were manually reviewed and refined across multiple rounds (Fig. 2A and B). This strategy progressively improved both label quality and model performance. We also applied class-weighted and morphology-sensitive loss functions during training to mitigate underrepresentation of rare or ambiguous structures. During inference, raw EM images were processed through two parallel branches (Fig. 2C): the MicroStructFormer (MSF) that generated comprehensive ultrastructure predictions (Fig. S1), and the AxonDeepSeg (ADS) that produced axon and myelin masks. The outputs of both branches were then merged to obtain a unified segmentation map that incorporates complementary strengths from each model (Fig. 2C).

**Fig. 2.**
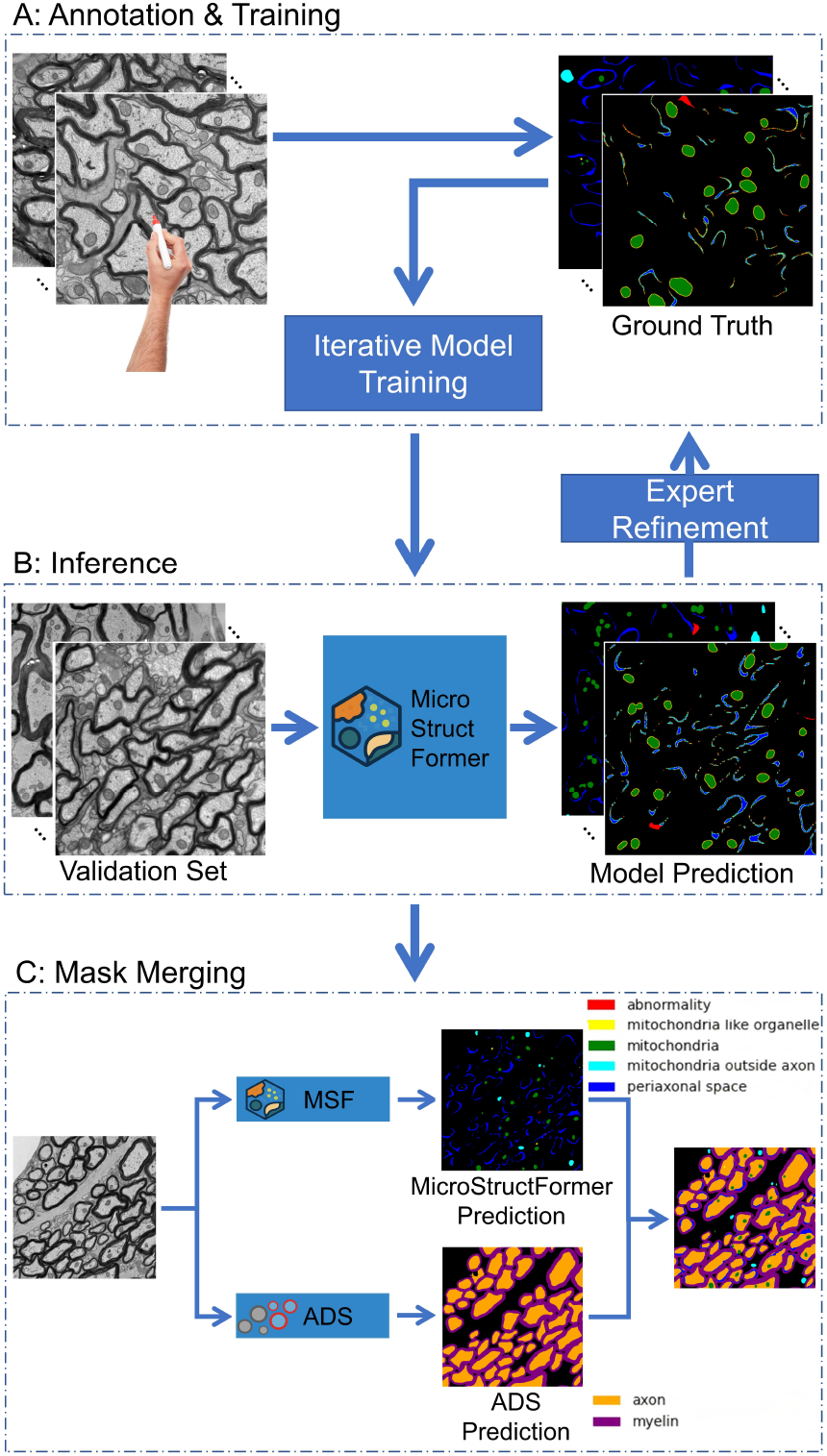
An overview of the three-step segmentation pipeline. **(A)** Workflow of dataset annotation and model training. Microstructures in the training set were manually annotated and curated by experts. **(B)** Inference of unlabeled datasets using trained MicroStructFormer (MSF) Model. **(C)** Integration of AxonDeepSeg (ADS) predictions for axon and myelin semantic segmentation with MicroStructFormer predictions for fine-grained ultrastructural components.

### Data Preprocessing

Because our training relied in part on publicly available annotated EM datasets, we first needed to standardize different input formats. The source data varied considerably, including (i) images stored in different formats (e.g., ImageJ-compatible stacks, PNG, JPG, or TIFF), (ii) ground-truth annotations provided either as JSON polygon files or as indexed PNG masks, and (iii) image sizes that were not uniform across datasets. To ensure compatibility with our training pipeline, we standardized all data through a unified preprocessing step.

In this step, raw images were converted to grayscale tensors and normalized, while masks were converted into categorical label maps. To increase robustness and avoid overfitting, we applied a series of augmentations, including D4 group transformations (rotations and flips) and non-rigid deformations (elastic transform, grid distortion), designed to reproduce common EM artifacts. We also applied photometric adjustments (brightness/contrast), and Gaussian noise to mimic EM background variability. Further, we implemented random crop-and-zoom to improve scale invariance. Each original image-mask pair was augmented into multiple variants, expanding the dataset and improving model robustness.

Finally, the preprocessed and augmented datasets were saved into a standardized directory structure (train, val, test) with consistent naming conventions, ensuring direct compatibility with the model training interface. The same preprocessing and augmentation procedures were also applied to our manually labeled datasets. These annotated datasets can be accessed from our Hugging Face repository. Final resizing, rescaling, and normalization for model input were handled separately at training and inference time by the model-specific image processor.

### Training configuration and computational environment

Training was performed on a high-performance computing cluster equipped with NVIDIA H100 GPUs under a CUDA 12.2 environment. During training, input images and their corresponding segmentation maps were processed using the Hugging Face `Mask2FormerImageProcessor` with proper config settings, which automatically resized all images to 384 × 384 pixels, rescaled pixel values by a factor of 1/255, and normalized them using the default channel-wise mean and standard deviation parameters. As a result, all samples were presented to the network at a standardized input resolution, ensuring a fixed tensor size during training and inference. Optimization was carried out using AdamW (47) with a base learning rate of 4 × 10−4 and a polynomial decay scheduler (48). The batch size was set to 4 for both training and evaluation. To ensure reproducibility, the random seed was fixed throughout training.

### Model inference

After training, the model was deployed on a local workstation for inference. Local evaluation was performed on a machine equipped with an Intel Core i5-9400 CPU (2.90 GHz), 16 GB RAM, and an NVIDIA GeForce GTX 1660 Ti GPU with 6 GB VRAM. The same preprocessing pipeline was applied during inference, such that each input image was resized to 384 × 384 pixels before being passed to the network. This ensured that inference was carried out at the same fixed input resolution as used during training.

### Metrics for assessment of semantic segmentation

For semantic segmentation, our model was trained using the standard composite loss function proposed in Mask2Former (49), which integrates three components: a cross-entropy loss (*L*_ce_) for object query classification, a binary cross-entropy loss (*L*_mask_) for mask prediction, and a dice loss (*L*_dice_) to enhance mask boundary accuracy. The resulting total weighted loss is given by

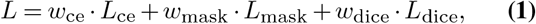

where *w*_ce_, *w*_mask_ and *w*_dice_ denote the corresponding loss weights, and *L*_dice_ is defined by

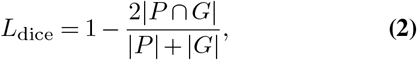

with *P* and *G* representing the predicted and ground truth masks, respectively.

For validation, model performance was assessed using the mean Intersection-over-Union (mIoU) metric, given by

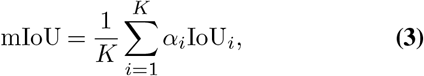

where

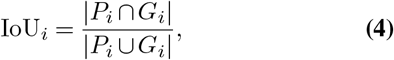

*K* is the number of evaluated classes, *α*_*i*_ is the weighting co-efficient for the i-th category, and *P*_*i*_ and *G*_*i*_ represent the predicted and ground truth masks for class *i*. During the whole process, certain parameters (such as class weights *L*_ce_ and the mIoU weighting coefficients *α*_*i*_) were modulated according to learning difficulty or rarity of specific instances, ensuring robust optimization across diverse cases. This protocol ensures a balanced assessment of both category recognition and pixel-level segmentation quality. Comparing the distributions of ground truth (GT) annotations and model predictions via split violin plots (Fig. S2), we found that while the model accurately captured overall genotype-dependent trends, a slight divergence was observed in the abnormality class, particularly in the most severe: double and triple KO conditions. This reduced sensitivity likely stems from the high morphological heterogeneity in myelin related pathological structures in these backgrounds, which increase the intrinsic difficulty of consistent segmentation. Nevertheless, the model effectively preserved the relative scaling of abnormalities across the investigated genotypes. For further details regarding the model, please refer to (49).

### Post-processing pipeline for axon–myelin morphometrics

#### Individual-level morphometrics from semantic masks

Pixel-wise EM segmentation alone does not generate fiber-level measurements, nor does it capture the spatial relationship between axons, myelin, mitochondria, periaxonal space and fiber abnormalities. To obtain biologically meaningful, axon-centered morphometrics, we converted semantic class masks into coherent fiber individuals. This was achieved by separating adjacent axons, assigning surrounding compartments to their corresponding fibers, and linking mitochondria to their most likely parent axon, thereby turning raw pixel labels into structurally consistent, fiber-centric representations.

Starting from the semantic class masks, we applied light morphological smoothing and then converted the pixel-wise labels into fiber-centered individuals through three sequential steps.

##### (1) Myelin-guided watershed

To separate adjacent axons into distinct axon instances, we slightly dilated the myelin mask to create a barrier between neighboring fibers, and used seeds derived from distance-transform maxima within each axon core to drive a marker-controlled watershed, thereby splitting touching axons into individual instances.

##### (2) Geodesic assignment of myelin, periaxonal space and abnormalities

To assign the surrounding intramyelin compartments to the correct axon, we performed geodesic partitioning of myelin, periaxonal space, and abnormalities within the region enclosed by each fiber. Before propagation, any axon interior unintentionally fragmented during semantic segmentation was merged so that each axon was represented by a single coherent core. We then initialized axon-specific seeds along the boundary of each axon core and propagated them outward over the geodesic distance field, allowing periaxonal space, fiber abnormalities, and any contacting or overlapping myelin to be assigned to the nearest axon in a geometry-aware manner. To avoid systematic under-assignment in large fibers, this propagation incorporated a mild radius-dependent scaling that preserved more uniform circumferential coverage across axons of different calibers.

##### (3) Mitochondria assignment

To associate mitochondria with their most likely parent axon, we assigned each mitochondrion component to the axon with which it shared the greatest boundary contact after slight dilation. When no direct contact was present, assignment was allowed only if the mitochondrion lay within a small tolerance zone just inside the axonal boundary, thereby reducing the risk of including extraxonal structures.

These steps provided per-fiber instances together with their associated myelin, periaxonal space, fiber abnormality, and mitochondrial regions.

Through these steps, semantic segmentation masks were converted into axon-centered instances (Fig. 3). Class-level predictions were first refined into distinct axon instances using a myelin-guided watershed (Fig. 3B). Myelin, periaxonal space, and abnormal regions of fiber were then geodesically assigned to their corresponding axons (Figs. 3C-E), and mitochondria were linked based on boundary contact or nearest-distance rules (Fig. 3F). This instance-level representation enabled consistent, fiber-centric morphometrics.

**Fig. 3.**
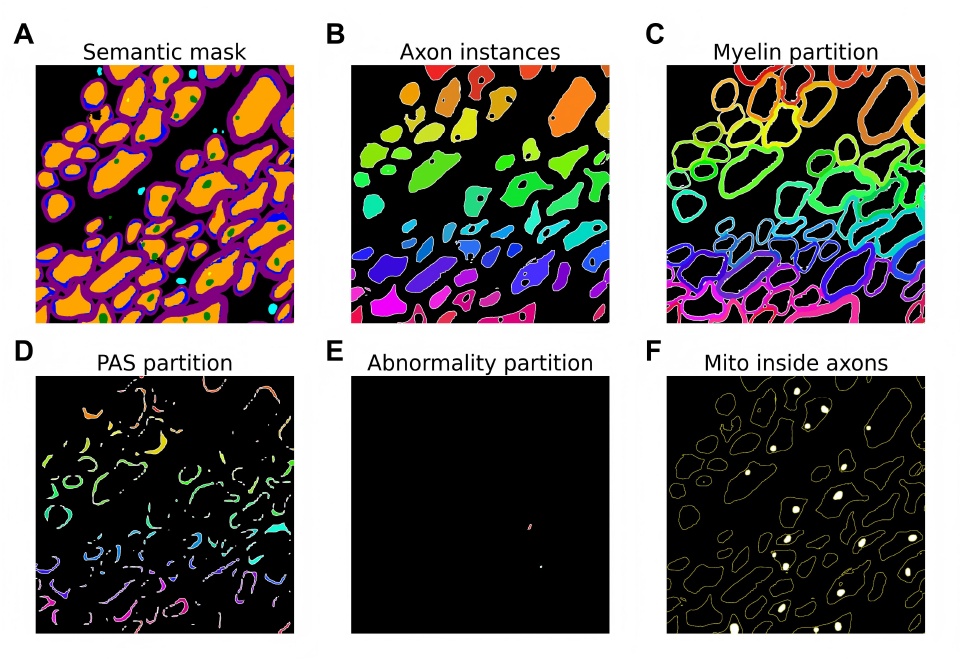
Fiber-specific decomposition of semantic segmentation masks. **(A)** Semantic segmentation mask showing class-level predictions for axon, myelin, mitochondria, PAS, and abnormal regions. **(B)** Axon instances generated using myelin-guided watershed segmentation. **(C)** Myelin partitioned and assigned to each axon based on geodesic proximity. **(D)** PAS partitioned per axon using geodesic grow-and-assign rules. **(E)** Abnormal regions assigned to their corresponding axons. **(F)** Instance-level mitochondria assignment, where each mitochondrion is linked to the axon with which it shares maximal boundary contact or nearest proximity. Together, these steps convert semantic masks into coherent axon-centered instances suitable for morphometric analysis.

#### Measurements

For each axon, we defined two fiber configurations for morphometric computation: (a) the exclusive fiber (used for all primary analyses) which includes only the axon and its associated compact myelin, and (b) the inclusive fiber that additionally incorporates the periaxonal space (PAS) and abnormal regions of uncompacted oligodendrocyte membrane, providing a broader anatomical context when needed. Axon and fiber (i.e., axon plus myelin) diameters: *D*_*ax*_ and *D*_*fib*_, respectively, were computed using a circular-equivalent diameter formulation

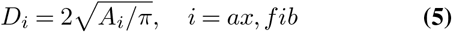

where *A*_*i*_ is the cross-sectional area of axon (*i* = *ax*) or fiber (*i* = *f ib*), and the fiber area satisfies *A*_*fib*_ = *A*_*ax*_ + *A*_*my*_, where *A*_*my*_ denotes the myelin cross-sectional area.

Based on the fiber individual we obtained, we computed the following morphometric ratios

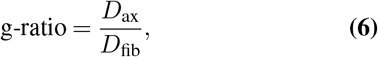

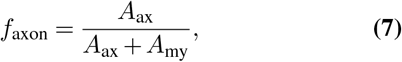

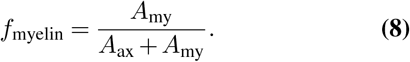

Because g-ratio is defined using the geometry of the entire axonal compartment, any intra-axonal structures (including mitochondria and mitochondria-like organelles) were retained as part of the axon area rather than subtracted. Per-image scale (µ m/pixel) is read from a calibration table and applied where available. To obtain image-level summaries with balanced contribution, we performed a simple per-image bootstrap, sampling 2,000 axon instances per image (optionally stratified by the presence of mitochondria), aggregating the results, and repeating this procedure across replicates to estimate g-ratio, myelin fractions, and mitochondria-myelin distances. The implementation relied on standard `scikit-image, scipy.ndimage` operators and fixed random seeds for reproducibility.

### Image-level morphometrics from semantic masks

For each image, instance-level measurements obtained from the segmentation pipeline were first filtered to exclude extremely small objects. Morphometric descriptors were then extracted for each semantic class (axon, myelin, PAS, abnormality, mitochondria, etc.). The full list of class-specific metrics and their definitions are provided in Supplementary Section A.2. Image-level features were generated by aggregating these individual-level measurements within each image-class combination. For every metric, a set of summary statistics was computed and stored in long format A.2, then reshaped into a wide table indexed by `ImageID, GenoType, Age` and `Region.` Genotype labels were harmonized using rule-based consolidation, and categorical variables (age, region, genotype) were one-hot encoded for downstream statistical analyses (e.g., violin plots and random forest classification).

### Statistical testing and multiple comparison correction

Statistical comparisons were performed using two-sided Mann-Whitney U tests. P-values were adjusted for multiple comparisons using the Holm-Bonferroni method. Unless otherwise specified, comparisons were made against the WT group within each analysis. Significance was summarized using standard notation (n.s., *, **, ***), corresponding to adjusted p-values.

### Descriptive genotype-level trend fitting

For Fig. 9E, we focused exclusively on myelinated fibers with complete metadata. We excluded axons touching the image border to prevent contour truncation, and removed those with an equivalent circular diameter below 0.5 µm. Fibers from CST, VM, and GC were pooled. For each genotype and age, we plotted the average mitochondrial area normalized to total axon area, estimating uncertainty through adaptive fiber-level bootstrap resampling.

We fitted the P30 and P90 mean estimates separately using a constrained decreasing Hill function where *x* represents the relative *Mbp*/*Gapdh* expression level and *y* represents the mean normalized mitochondrial area fraction,

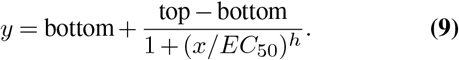

To account for uncertainty, we weighted each fit using one-half of the bootstrap 95% confidence-interval width. The fits were constrained to non-negative values for bottom, *EC*_50_, and the amplitude term (top − bottom), while restricting the Hill coefficient to 0.01 ≤ *h* ≤ 10. We did not estimate parameter confidence intervals. Because the bootstrap intervals characterize genotype-level mean uncertainty rather than the variation of the fit parameters, we interpreted the resulting curves as descriptive summaries instead of formal parameter inference. Since the fitting weights were derived from bootstrap intervals obtained by resampling fibers after pooling across images, regions, and animals, they reflect fiber-level uncertainty in the genotype-level means.

### Partial correlation

To assess the association between mitochondrial area and g-ratio independently of fiber diameter, we computed partial correlations while controlling for fiber diameter. The latter provides a more integrative descriptor of axon-myelin geometry than axon diameter alone and has been shown to better capture g-ratio scaling relationships across fiber populations (50).

Because the relationships between morphometric variables often follow nonlinear scaling laws, variables were first log-transformed to linearize their relationships and stabilize variance across the dynamic range (51). Partial correlation 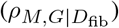 was then quantified as the Spearman correlation between the residuals obtained by regressing the mitochondrial area and g-ratio on the log-transformed fiber diameter, as follows

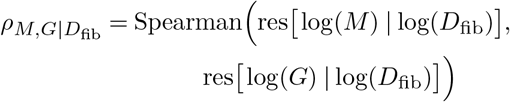

where *M* = mitochondrial area and *G* = g-ratio.

### Data and code availability

Annotated EM datasets, trained model weights, analysis code, documentation, and usage examples associated with this study are publicly available through the project website: https://dnq2023.github.io/MSF_WEBSITE/. These resources are provided to support reproduction of the reported analyses and reuse of the MicroStructFormer workflow.

## Results

### Quantitative analysis per image

#### Per-image ultrastructure area analysis

Pronounced differences in myelin-related ultrastructural features are associated with diminished *Mbp* mRNA levels. To systematically quantify these image-level changes, we applied our automated multi-compartment EM segmentation pipeline followed by pooled statistical analysis across genotypes to evaluate how area fractions of myelin, abnormalities, mitochondria and periaxonal space change (Fig. 4, Figs. S3–S12).

**Fig. 4.**
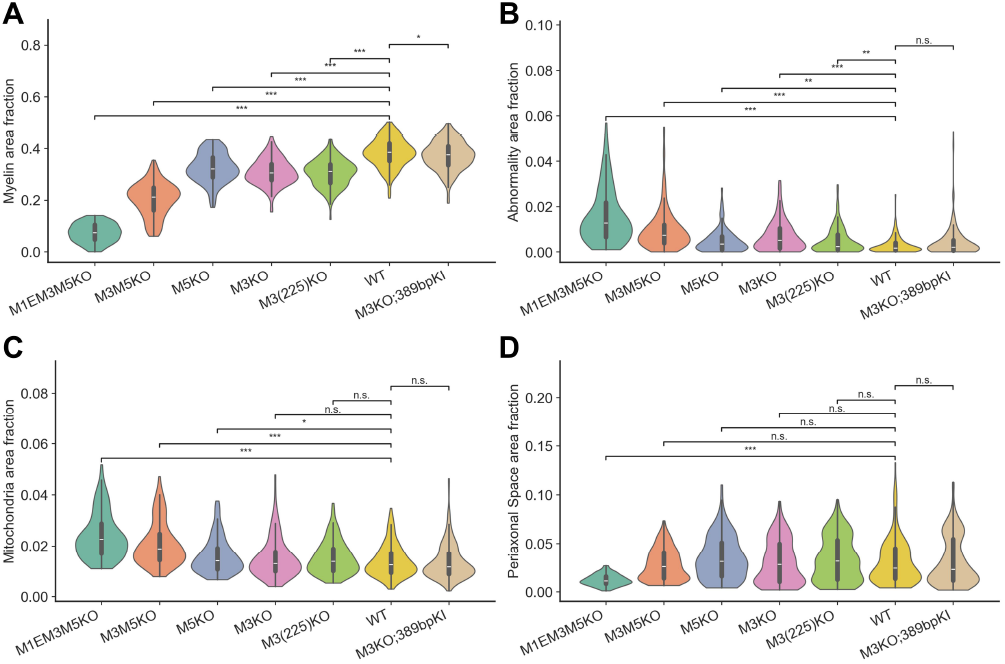
Distributions of ultrastructural features across *Mbp* genotypes pooled across ages and regions. Violin plots showing pooled per-image distributions of area fraction of (A) myelin, (B) abnormality, (C) mitochondria, and (D) PAS across different *Mbp* genotypes. Notice how a reduction in *Mbp* mRNA expression leads to an increase in mitochondrial area fraction representing higher metabolic demand. Significance was assessed using two-sided Mann-Whitney U tests with Holm-Bonferroni correction; asterisks denote adjusted p-values (* p < 0.05, ** p < 0.01, *** p < 0.001, Holm-Bonferroni corrected).

A progressive decline in myelin thickness was observed alongside reduced *Mbp* expression, in line with the established structural role of MBP in forming the major dense line of compact myelin, with the most pronounced reduction observed in double and triple KO groups (Fig. 4A). In contrast, the prevalence of abnormal structures increases as *Mbp* expression decreases (Fig. 4B), which may reflect the presence of non-compacted oligodendrocyte plasma membrane or ongoing remodeling processes under low-*Mbp* conditions (Fig. S13). In parallel, mitochondrial content within axons also increases under *Mbp* deficiency (Fig. 4C, Fig. S14), supporting the idea that impaired myelin insulation imposes an elevated metabolic burden to maintain efficient axonal conduction.

Consistent with an inverse relationship, we also found that myelin deficiency is associated with an increase in mitochondrial number (Fig. S11). This indirectly supports the notion that, even though previous studies (30) focused on Shiverer mice, while our work addresses *Mbp*-deficient hypomyelination, the physiological compensatory mechanisms appear to be similar in both cases.

To characterize further changes in axonal composition, we quantified the fraction of unmyelinated axon area across genotypes, ages, and regions (Fig. S7). Across conditions, unmyelinated axon area is progressively increased in KO groups, particularly under more severe *Mbp* perturbations. The resulting compositional shift indicates a substantial reorganization of axonal populations, toward fibers with reduced or absent myelin wrapping. Such compositional changes provide an important structural context for subsequent analyses, because shifts in the myelinated-versus-unmyelinated axon balance can influence both mitochondrial and myelin-related image-level metrics.

Interestingly, our quantitative analysis also shows that changes in periaxonal space exhibit a biphasic pattern, with a modest increase in single KO (potentially reflecting enhanced axon-oligodendrocyte interactions), followed by a marked decline in double and triple KO conditions. We interpret this decrease not as a true loss of periaxonal space, but as a consequence of severe morphological disruption, in which the canonical organization of the periaxonal space is replaced or obscured by abnormal and entangled structures (Fig. S13). Furthermore, at the image level, mitochondrial, axonal, and myelin area fractions exhibit positive associations across genotypes (Fig. S15), compatible with geometric scaling effects and differences in the relative area of axons across images. However, these population-level correlations do not resolve whether such relationships persist at the level of individual fibers or reflect underlying structural dependencies.

To further contextualize the pooled violin trends of ultrastructural features across Mbp genotypes (Fig. 4), we summarized each genotype condition by its mean area fraction and plotted mitochondrial (Fig. 5) and myelin area fractions (Fig. 6) against relative *Mbp*/*Gapdh* expression across regions and ages. For each genotype, mitochondrial area fraction shows a consistent inverse monotonic trend with relative *Mbp*/*Gapdh* in all six panels (Spearman ρ *<* 0 in each region-age sub-set), whereas myelin area fraction shows a corresponding positive monotonic trend (Spearman ρ > 0 in all six panels). Although these relationships visually appear close to being linear across most panels, we interpret them conservatively as monotonic genotype-level associations. It’s important to note that myelin area fraction shows relatively limited genotype-level dispersion around the fitted trend, consistent with the direct role of *Mbp* in myelin formation and maintenance. In contrast, mitochondrial area fraction shows a little bit more region- and age-dependent variability, with narrower genotype-level spread at P90 than at P30, and generally smaller spread in VM across ages.

**Fig. 5.**
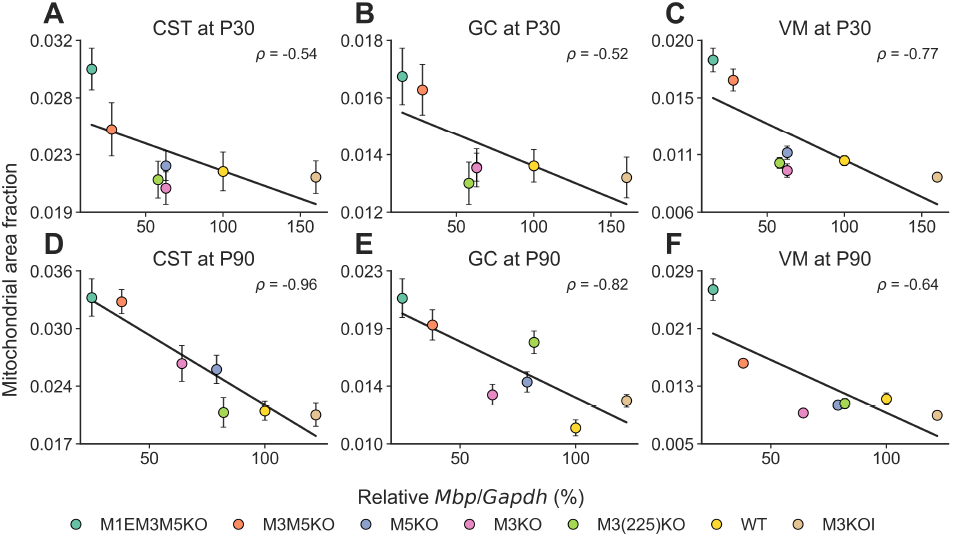
Genotype-level association between relative *Mbp*/*Gapdh* expression and mitochondrial area fraction. Scatter plots showing mitochondrial area fraction versus relative *Mbp*/*Gapdh* expression across *Mbp* genotypes in three spinal cord regions: (A,D) CST, (B,E) GC, and (C,F) VM, at P30 (A-C) and P90 (D-F). Each point represents one genotype condition, and error bars indicate variability across images within each condition. Black lines indicate linear fits, and ρ values denote Spearman correlations. Error bars are shown where visible; when not apparent, they are smaller than or partially obscured by the plotted symbols.

**Fig. 6.**
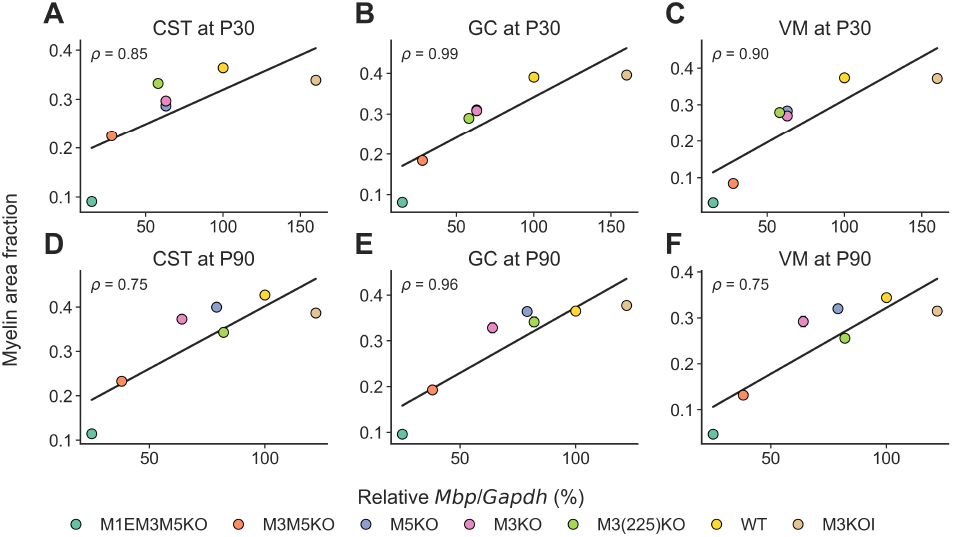
Genotype-level association between relative *Mbp*/*Gapdh* expression and myelin area fraction. Scatter plots showing myelin area fraction versus relative *Mbp*/*Gapdh* expression across *Mbp* genotypes in three spinal cord regions: (A,D) CST, (B,E) GC, and (C,F) VM at P30 (A-C) and P90 (D-F). Each point represents one genotype condition, and error bars indicate variability across images within each condition. Black lines indicate linear fits, and ρ values denote Spearman correlations. Error bars are shown where visible; when not apparent, they are smaller than or partially obscured by the plotted symbols.

#### Random forest prediction of genotype

To determine whether fiber ultrastructural phenotypes alone are sufficient to distinguish between *Mbp* enhancer-edited mouse lines, we trained a random forest classifier on a comprehensive set of axon-, myelin-, PAS-, abnormality- and mitochondria-derived morphometrics (Fig. 7A). This analysis was performed on the pooled dataset across all available ages (P30, P90, P400 where available) and all three spinal cord regions (CST, GC, VM). Region were included as one-hot encoded covariates in the same model, whereas age was not included to avoid label leakage from missing P400 data in some genotypes.

**Fig. 7.**
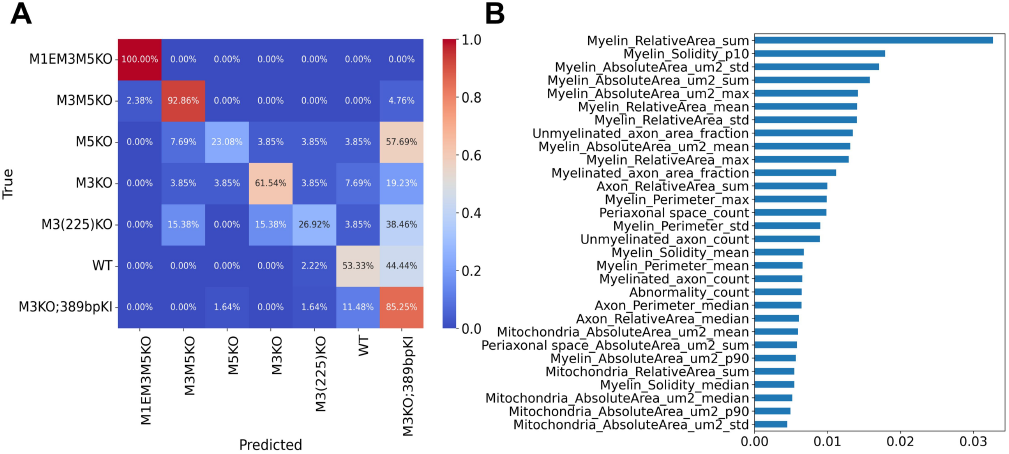
Random forest classification of genotypes using ultrastructural features. **(A)** Confusion matrix of genotype classification. Severe enhancer deletions are well separated (e.g., M1EM3M5KO, 100%; M3M5KO high accuracy), with errors largely confined within this subgroup, whereas milder perturbations show greater overlap and are frequently assigned to M3KOKI. **(B)** Top 30 features ranked by mean decrease in impurity (MDI). Myelin-related metrics dominate, followed by axonal geometry and mitochondrial features.

Genotypes associated with severe enhancer deletions exhibited near-perfect separability. In particular, M1EM3M5KO is classified with 100% accuracy, and M3M5KO shows similarly high within-class accuracy with minimal misclassification. When misclassification occurred among these severe genotypes, it is largely confined within this severely perturbed subgroup, with very limited confusion with milder perturbations or WT, indicating, as expected, that large-scale *Mbp* disruption produces highly distinct and internally consistent ultrastructural signatures.

In contrast, genotypes with milder perturbations display greater overlap. Single KO and WT conditions are more frequently misclassified, with a substantial fraction of these samples being assigned to the M3KOKI class. Rather than reflecting random error, this pattern suggests a structured and biologically adjacent relationship in feature space. Indeed, the M3KOKI line undergoes an initial *Mbp* deletion followed by rescue, and may therefore retain residual ultrastructural features associated with earlier knockout states while partially recovering to or beyond WT morphology. In addition, transient or hypermyelination-related remodeling with age in this line may further broaden its feature distribution, potentially explaining why most M3KOKI samples remain correctly assigned while a subset of WT samples is also mapped to this class. As a result, this group is likely to span a broader morphological spectrum that overlaps with both single KO and WT phenotypes. This broadened distribution likely causes the M3KOKI class to occupy a nearby or partially overlapping region of feature space; namely, when the classifier encounters intermediate or ambiguous phenotypes, they are preferentially assigned to this group. In agreement with this interpretation, misclassifications are not uniformly distributed but instead exhibit a directional bias toward the M3KOKI class.

Conducting feature importance analysis further revealed that myelin-based metrics are the most dominant predictors of genotype (Fig. 7B). Feature importance was quantified using the random forest mean decrease in impurity (MDI), defined as the average reduction in node impurity contributed by a feature across all trees in the ensemble. Relative myelin area (both mean and sum), together with solidity and perimeter statistics, showed the largest contributions, followed by axon caliber-related measures and a smaller subset of mitochondrial, PAS, and abnormality features. In this pooled model, region shows comparatively low feature importance relative to the dominant descriptors.

Although mitochondria-, PAS-, and abnormality-derived metrics contribute less than the principal myelin descriptors, their consistent presence among the informative features indicates that organelle content, periaxonal geometry, and localized structural defects provide additional, albeit weaker, discriminatory signals. Thus these classification and feature importance analyses support the notion that quantitative ultra-structural signatures encode robust, yet hierarchically structured, genotype-specific information, with strong separability for severe perturbations and graded overlap among milder conditions. Consistent trends were observed when restricting the analysis to features derived from individual compartments, including axon-, myelin-, mitochondrial-, and periaxonal space-specific subsets (Fig. S16, Fig. S17, Fig. S18, and Fig. S19)

### Quantitative analysis per individual fiber

#### Per-fiber ultrastructure area analysis

While the per-image summaries reveal clear group-level shifts, they average over heterogeneous populations of fibers within each image. As a result, they cannot disentangle how key structural variables (such as fiber diameter, mitochondrial content, and myelin thickness) relate to each other within individual axons. These population-level analyses therefore provide important context, but lack the resolution required to resolve fiber-specific relationships.

We therefore performed per individual fiber analyses, treating each axon and its associated compartments as individual data points. This allowed us to determine whether the relationships observed across genotypes are preserved at the level of individual fibers, before explicitly accounting for axonal geometry in subsequent analyses.

To further resolve the heterogeneity underlying image-level trends, we examined ultrastructural features at the level of each fiber across genotypes (Fig. 8A). As expected (28), the quantification of g-ratio distributions (Fig. 8B) revealed a progressive rightward shift with increasing severity of *Mbp* deletion, consistent with reduced myelin thickness.

**Fig. 8.**
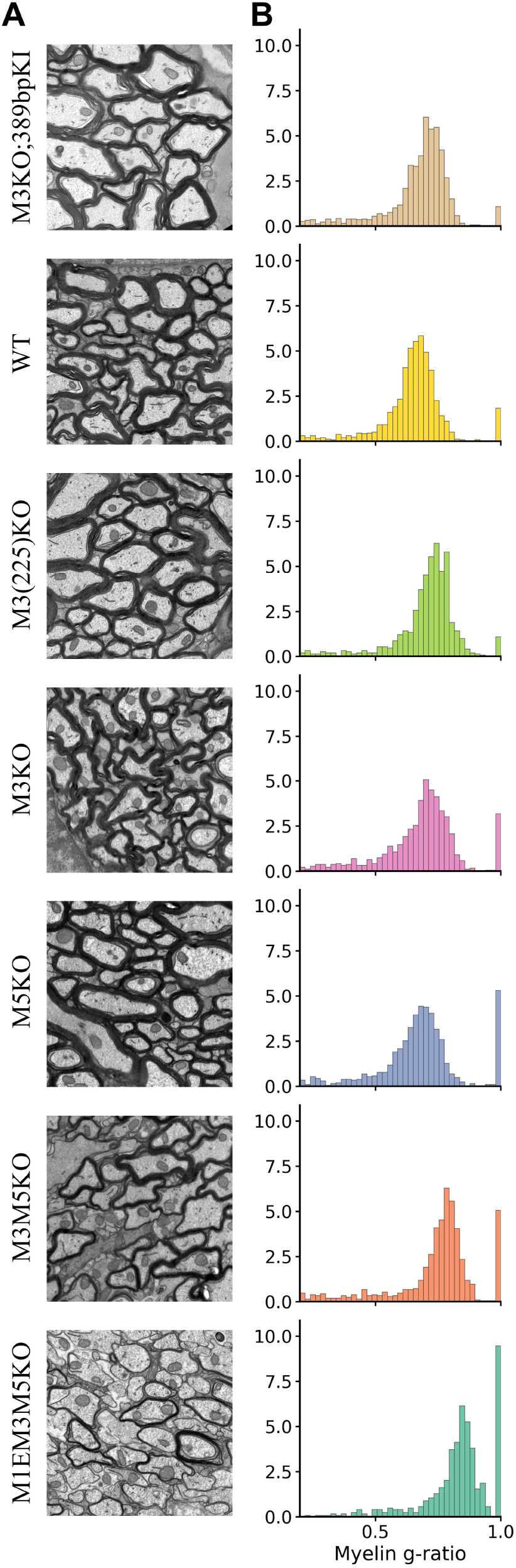
Raw ultrastructure and myelin g-ratio across genotypes (P90 CST). **(A)** Representative electron microscopy images from P90 CST, displayed for each genotype. **(B)** Distribution of myelin g-ratio values, calculated from circle-equivalent diameters derived from axon and myelin areas, with periaxonal space and abnormal regions omitted from the outer boundary definition.

We next asked whether mitochondrial changes observed at the image level are similarly preserved at the level of individual fibers. To determine this, we normalized per-axon mitochondrial area to the WT mean within each myelination state (Fig. 9A, B). In agreement with the image-level observations, elevated mitochondrial area is most apparent in the more severe KO groups (double and triple KO conditions), and these differences remain statistically significant at the per-axon level. This increase is most clearly preserved in myelinated fibers, whereas unmyelinated fibers show weaker shifts and broader overlap with WT. In contrast, single KO conditions do not exhibit a consistent or significant increase. Together with the increased mitochondrial number observed at the image level (Fig. S11), the persistence of this per-fiber increase indicates that the image-level trend is not driven solely by differences in mitochondrial number or overall tissue composition across genotypes, but also by changes in mitochondrial area itself.

**Fig. 9.**
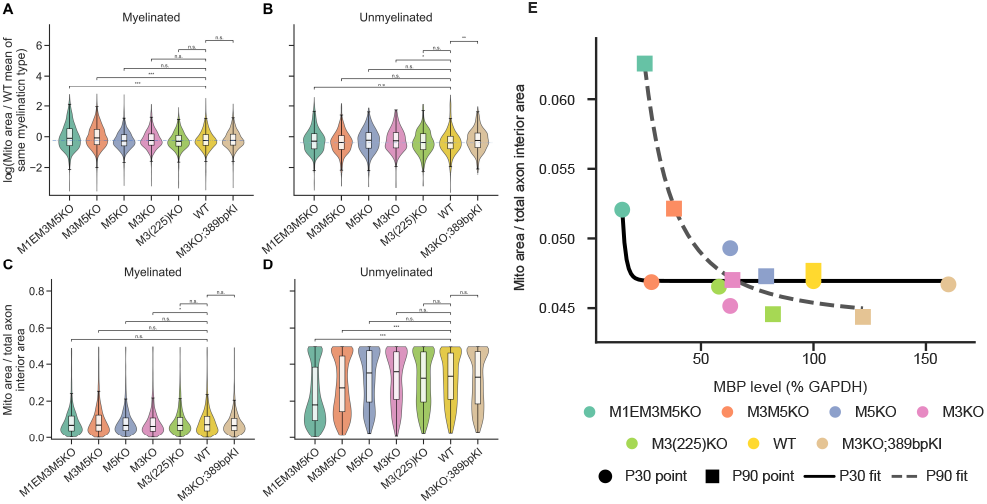
Per-fiber mitochondrial area and mitochondrial burden across *Mbp* genotypes, pooled across ages and regions. **(A, B)** Violin plots showing per-axon mitochondrial area normalized to the WT mean of the corresponding myelination state, pooled across all analyzed spinal cord regions and ages and separated into myelinated **(A)** and unmyelinated **(B)** populations across *Mbp* genotypes, with values displayed on a log scale. Black dots and error bars indicate group means and associated uncertainty. Horizontal dashed lines denote the WT median for myelinated and unmyelinated axons. Significant increases in absolute mitochondrial area are primarily restricted to the more severe KO groups in myelinated fiber (double and triple KO conditions), whereas single KO conditions largely overlap with WT. **(C, D)** Violin plots showing mitochondrial area normalized by total axon interior area for mitochondria-positive axons, separated into myelinated **(C)** and unmyelinated **(D)** populations across *Mbp* genotypes. Boxplots indicate the median and interquartile range. In myelinated axons, mitochondrial burden remains relatively stable across genotypes, whereas unmyelinated axons show reduced and more variable mitochondrial burden under increasing *Mbp* perturbation (* p < 0.05, ** p < 0.01, *** p < 0.001, Holm-Bonferroni corrected for panels A-D). **(E)** Scatter plot of all-region pooled genotype-level per-fiber arithmetic means for mitochondrial area normalized by total axon interior area, plotted against relative *Mbp*/*Gapdh* level (% GAPDH) at P30 and P90. Each point represents one genotype-age condition; colors encode genotype and marker shapes encode age. Solid and dashed curves show descriptive constrained decreasing Hill-function fits for P30 and P90, respectively, weighted by bootstrap mean uncertainty. These fits summarize trends and are not used for parameter-level inference. Descriptive Hill fit parameters for P30 and P90 are provided in Table 1.

These results show that mitochondrial expansion is not only a consequence of overall tissue composition, but a robust phenotype preserved at the resolution of individual fibers. Notably, this phenotype is not distributed uniformly across axonal states: in absolute terms, mitochondrial accumulation is preferentially evident in myelinated fibers, suggesting that severe *Mbp* deficiency is associated with selective preservation or accumulation of mitochondrial content within the myelinated population. Furthermore, the threshold-like behavior observed here, in which significant mitochondrial accumulation emerges only under severe *Mbp* deficiency, is consistent with a severity-dependent compensatory response or a stress-associated remodeling under advanced demyelination.

Importantly, this analysis (Fig. 9A, B) still reflects absolute mitochondrial content at the per-fiber level, expressed relative to the WT mean of the corresponding myelination state, rather than burden normalized by axon interior area. When this relationship is examined by normalizing mitochondrial area to axon interior area, distinct patterns emerge depending on myelination state: myelinated axons show relatively stable mitochondrial burden across genotypes, whereas unmyelinated axons exhibit reduced and more variable mitochondrial allocation under increasing *Mbp* perturbation (Fig. 9C, D). To further examine this relationship, we evaluated normalized mitochondrial burden against relative *Mbp*/*Gapdh* levels in the myelinated-fiber subset after pooling CST, VM, and GC (Fig. 9E). Constrained decreasing Hill curves were fitted independently to the P30 and P90 genotype-level mean estimates. The P90 series showed a clearer negative pattern as expression increased, whereas the P30 series declined at the low-expression end and then remained close to a plateau. These curves summarize genotype-level bootstrap mean estimates and are intended primarily to illustrate descriptive age-specific trends.

Taken together, the comparison between absolute and normalized measures reveals an informative reversal in interpretation. In myelinated fibers, mitochondrial area increases under severe *Mbp* deficiency, yet normalization by axon interior area reduces or eliminates these differences, indicating that the absolute increase largely scales with the size of the axon interior rather than reflecting a disproportionate increase in mitochondrial burden. In unmyelinated axons, by contrast, absolute mitochondrial area shows only limited differences across genotypes, whereas mitochondrial burden tends to decrease after normalization, indicating that these fibers do not preserve mitochondrial allocation relative to axon interior area under severe perturbation. This divergence indicates that mitochondrial expansion is not uniformly distributed across axons, but instead reflects a selective redistribution of mitochondrial content that is preferentially maintained in myelinated fibers.

#### Partial correlation

Within myelinated fibers, we next performed partial correlation analyses to assess the relationship between mitochondrial content and g-ratio after controlling for fiber diameter (Fig. 10). As shown previously, fiber diameter represents a major structural variable influencing both mitochondrial abundance and myelin thickness, introducing a potential geometric confound in their apparent association (Fig. S20, S21).

**Fig. 10.**
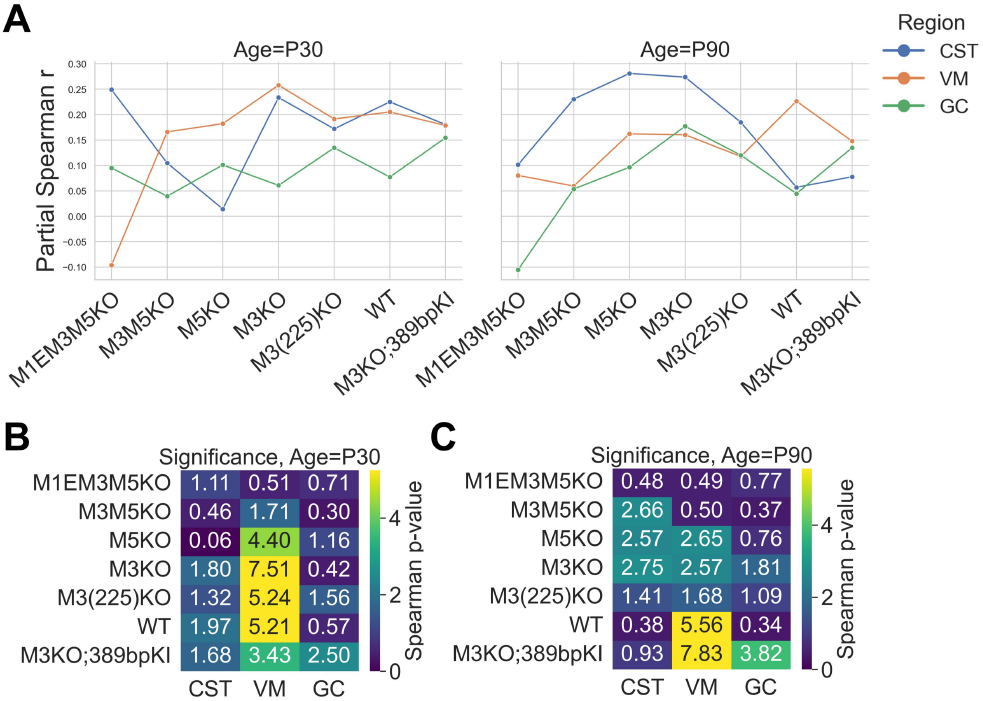
Partial correlation between mitochondrial content and g-ratio controlling for fiber diameter. **(A)** Partial Spearman correlation coefficients between mitochondrial area and g-ratio across genotypes, stratified by age and spinal cord region. **(B, C)** Significance levels (− log_10_(*p*)) associated with the partial correlations shown in **A**, for P30 **(B)** and P90 **(C)**.

To remove this effect, we first regressed log-transformed mitochondrial metrics and g-ratio against log-transformed fiber diameter. Residuals from these models were then used to quantify diameter-independent associations using Spearman correlation within each analyzed age, genotype, and region. A summary of the partial correlation coefficients across genotypes for each region at P30 and P90 were then obtained (Fig. 10A). Examining the corresponding significance maps for P30 (Fig. 10B) and P90 (Fig. 10C) reveals that despite clear genotype-dependent shifts in both mitochondrial area and g-ratio at the population level, diameter-controlled partial correlations within individual groups are generally weak and frequently close to zero. This indicates that much of the apparent mitochondria-g-ratio association arises from shared dependence on fiber diameter and genotype-dependent shifts in overall mitochondrial and myelin levels, rather than from a stable fiber-level coupling between mitochondrial abundance and myelin thickness.

In particular, although mitochondrial content is elevated in severe *Mbp* KO conditions, this increase does not translate into a consistent relationship with myelin thinning across individual fibers within the same group. Instead, mitochondrial remodeling appears to reflect coordinated responses within specific biological groups, rather than a fine-grained, fiber-by-fiber scaling with myelin structure.

Although there is an overall lack of stable coupling, several context-dependent patterns can be observed. In specific region-age combinations (most prominently CST at P90), partial correlations exhibit a non-monotonic trend across genotypes, increasing in intermediate KO conditions and decreasing in more severe perturbations (Fig. 10A), in line with trends observed in earlier image-level and per-fiber analyses (Fig. S22, Fig. S23). In addition, the WT group shows relatively higher partial correlations at early developmental stages (P30), which diminish at later developmental stages (Fig. 10A), potentially reflecting more coordinated developmental scaling during early maturation that becomes stabilized in adult tissue. Temporal trends in Fig. S24 and Fig. S25 provide additional developmental context, with mitochondrial area fraction in KO lines tending to peak at intermediate stages, whereas myelin area fraction continues to increase with age. However, these patterns are not uniformly observed across regions and conditions and should therefore be interpreted as contingent on specific biological constraints rather than indicative of a consistent underlying coupling.

## Discussion

To characterize genotype-dependent shifts in ultrastructural composition at the image level, we first quantified per-image area fractions of major axonal and myelin-associated ultrastructures. Our results showed that myelin content decreases progressively with reduced *Mbp* expression (Fig. 4), consistent with the established role of MBP in forming the major dense line of compact myelin, with the most pronounced changes observed in double and triple KO groups. Abnormal structures increase as *Mbp* expression decreases, which we interpret as reflecting either the presence of non-compact myelin or ongoing oligodendrocyte-mediated remodeling processes in low-*Mbp* conditions (Fig. S13). Mitochondrial content within axons also increases under *Mbp* deficiency, matching prior observations in dysmyelinated CNS, including *shiverer* mice (30, 31). Such mitochondrial accumulation has been associated with altered axonal metabolic demands under impaired myelin insulation (44).

While periaxonal space exhibits a biphasic pattern, with a modest increase in single KO conditions followed by a marked decline in double and triple KO conditions, we interpret this decrease not as a true loss of periaxonal space, but as a consequence of severe morphological disruption, in which the canonical periaxonal architecture is disrupted or no longer clearly preserved (46, 52).

Building on these distributional patterns, image-level regressions revealed positive associations between mitochondrial content, axonal area, and myelin metrics across genotypes (Fig. S15). These trends are consistent with shared geometric scaling and variation in axonal area across images, whereby images enriched in larger or more numerous axons exhibit increased myelin and mitochondrial area (36, 53). However, the strength of these associations remained modest and variable across conditions, suggesting that these image-level correlations may be influenced by underlying structural composition and geometric effects, and motivating more granular analyses at the level of individual fibers.

When genotype conditions were explicitly arranged by relative *Mbp*/*Gapdh* expression (Figs. 5 and 6), the direction of the genotype-level trends became more explicit: mitochondrial area fraction increased as relative *Mbp*/*Gapdh* expression decreased, whereas myelin area fraction increased as relative *Mbp*/*Gapdh* expression increased. This ordering-based view complements Fig. 4 by showing that genotype effects were not only shifted in their pooled distributions, but also organized along a coherent condition-level expression gradient. Myelin area fraction showed relatively limited genotype-level dispersion, whereas mitochondrial area fraction showed structured tract- and age-dependent variability, with condition-level spread narrowing at P90 and remaining comparatively smaller in VM. These EM-derived spinal cord findings mirror recent MRI observations in the brain (54).

At the per-fiber level, we found that mitochondrial content is increased in severe KO conditions, in agreement with prior reports of elevated axonal mitochondrial activity and mitochondrial accumulation in dysmyelinated CNS axons, including the *shiverer* mouse (30, 55). This observation supports the conclusion that the image-level trends are not solely driven by tissue composition. However, when mitochondrial content is normalized by axonal area, distinct state-dependent patterns emerge, with myelinated axons maintaining relatively stable mitochondrial burden, whereas unmyelinated axons exhibit reduced and more variable mitochondrial burden under increasing *Mbp* perturbation (Fig. 9C, D). Evaluating the data by expression level further shows that while relative *Mbp*/*Gapdh* expression and normalized mitochondrial burden share an inverse relationship, the dynamics of this decline vary between P30 and P90 within myelinated fibers (Fig. 9E). In the pooled dataset, the P90 fit exhibited a broader dynamic range and a sustained decrease with increasing MBP proxy levels. Conversely, the P30 curve declined initially but quickly plateaued. This age-dependent divergence suggests that at P30, mitochondrial differences are restricted primarily to genotypes with severe expression loss, whereas by P90, they manifest across a broader genotypic spectrum. This aligns with a progressive, age-dependent accumulation of mitochondrial burden driven by chronic myelin-related metabolic stress (17, 30, 40, 56). However, because these curves are based on aggregated genotype-level estimates, they should be viewed as summaries of broad trends rather than evidence for a strict dose-response relationship. These findings indicate that mitochondrial changes at the level of individual fibers are not uniform across axonal states, and reveal a clear divergence between absolute and normalized measures: whereas mitochondrial area increases in myelinated fibers under severe *Mbp* deficiency, normalization by axonal area reduces or eliminates these differences, and in unmyelinated axons mitochondrial burden can even decrease. This pattern indicates that mitochondrial expansion is not uniformly distributed across axons, but instead reflects a selective redistribution of mitochondrial resources, preferentially maintained in myelinated fibers while unmyelinated axons show a relative deficit in mitochondrial allocation.

One possible interpretation of this divergence is that axons differ in their capacity to receive and sustain metabolic support under impaired myelination. In myelinated fibers, mitochondrial content may be relatively preserved through axon–oligodendrocyte metabolic coupling, whereas in unmyelinated axons such support is reduced or absent, leading to lower relative mitochondrial allocation (57, 58). A complementary possibility is that severe myelin disruption alters activity-dependent mitochondrial transport and localization, producing a non-uniform distribution of mitochondrial content across fibers rather than proportional scaling with axonal size (59–61). Although direct evidence for these mechanisms is beyond the scope of the present study, both possibilities are compatible with the heterogeneous single-fiber allocation pattern observed here.

Further analysis identified fiber diameter as a primary structural variable influencing both mitochondrial content and g-ratio. Bootstrap analyses additionally revealed variability in mitochondria–myelin relationships across genotypes (Fig. S22, Fig. S23), indicating that these associations are not stable. When analyses were restricted to myelinated fibers and fiber diameter was controlled using partial correlation, the mitochondria–g-ratio associations became largely inconspicuous (Fig. 10). Partial correlations are generally weak and inconsistent, indicating that much of the apparent relationship arises from shared geometric dependence on axonal caliber rather than a direct coupling between mitochondrial abundance and myelin thickness. This statistical decoupling supports a scale-dependent interpretation of mitochondrial remodeling under *Mbp* perturbation: genotype-level analyses reveal coordinated shifts in mitochondrial content and myelin structure, but these changes do not translate into a stable fiber-level relationship once axonal size is taken into account. Instead, mitochondrial remodeling may be better understood as a population-level adjustment in axonal metabolic state within a perturbed axon–myelin unit, rather than a precise, fiber-by-fiber scaling of mitochondrial burden relative to local myelin thickness (17, 56).

Within this framework, the residual mitochondria–g-ratio relationship appears to be conditional on specific biological contexts rather than reflecting a universal coupling rule. In particular, the non-monotonic trends observed across genotypes, with partial correlations increasing in intermediate perturbation states and decreasing in more severe KO conditions, are in line with an initially coordinated remodeling response that becomes less organized as *Mbp* deficiency progresses. This pattern may reflect a transient alignment between mitochondrial content and myelin disruption that breaks down under more severe structural perturbation.

This context-dependency is also not uniform across spinal tracts, suggesting that the structural differences described above may rest on distinct cellular and functional bases. In the CST at P30, stronger positive partial correlations are already observed in double- and triple-perturbation conditions, suggesting earlier or more pronounced coordination between mitochondrial remodeling and myelin alteration. A plausible explanation is that the CST, as a major descending motor pathway, operates under higher functional and metabolic constraints, necessitating earlier adaptation of axon–myelin interactions when myelin integrity is compromised (62). This interpretation is compatible with evidence that oligodendrocyte composition and maturation differ across spinal tracts, with distinct mature oligodendrocyte populations preferentially associated with sensory versus corticospinal pathways and showing age-dependent changes in their relative abundance (63, 64). It is also compatible with recent findings that oligodendrocytes dynamically couple axonal activity to metabolic support, suggesting that pathways with higher functional demand may exhibit earlier or stronger remodeling of axon–myelin metabolic relationships (65).

Taken together, these findings indicate that mitochondrial remodeling under *Mbp* perturbation is governed by hierarchical structural constraints. Global shifts in mitochondrial and myelin organization emerge as a coordinated population-level response, whereas diameter-independent coupling at the level of individual fibers is weak and context-dependent, varying with myelination state, developmental stage, and regional tract context. This suggests that individual axons do not strictly adhere to a universal scaling rule, but instead exhibit a flexible rather than hard-coded metabolic adaptation. A primary limitation of this study is the limited number of biological replicates available for ultrastructural analysis. Because each EM dataset required a terminal experiment and extensive downstream processing, only one or two animals were available for several genotype-age conditions. Therefore, the quantitative comparisons presented here should be interpreted as evidence of reproducible structural trends rather than definitive population-level estimates. Future studies with larger cohorts will be required to validate these findings and to enable statistical frameworks that more rigorously partition biological and technical sources of variability. Nevertheless, the consistent directionality observed across conditions supports the interpretation that the reported changes reflect non-random ultrastructural shifts.

Overall, our results support a scale-aware view of dysmyelinated axons: genotype-level remodeling of mitochondrial and myelin organization is evident, but stable diameter-independent mito–myelin coupling at the single-fiber level is limited. This framework provides a quantitative basis for future studies to test how metabolic adaptation and myelin disruption interact across developmental stage, tract context, and perturbation severity.

## ACKNOWLEDGEMENTS

This work was supported by the Natural Sciences and Engineering Research Council of Canada (NSERC) Discovery (RGPIN-2019-04520) and Alliance International (ALLRP 588367-23) grants to AK. ND was partially supported by the China Scholarship Council (CSC) Joint Scholarship Program and the UNIQUE Excellence Scholarship 2026-2027 (Doctoral level). This research was enabled in part by computational resources provided by Calcul Qubec (https://www.calculquebec.ca) and the Digital Research Alliance of Canada (https://alliancecan.ca/). The funders had no role in study design, data collection and analysis, decision to publish, or preparation of the manuscript.

