## Supplementary PDF for "Deep learning-based decoding of axonal ultrastructure in gene-edited mice using electron microscopy imaging"

### Supplement

#### A. Supplementary Information.

***A.1. Per-image regression.*** To examine whether coordinated shifts across ultrastructural compartments (Fig. 4) reflect systematic relationships between individual features, we assessed image-level correlations between mitochondrial, axonal, and myelin area fractions. Across all genotypes, image-level regressions showed positive associations between mitochondrial and myelin area fractions (Fig. S15A).
Slopes were consistently positive, although the explained variance remained modest for mitochondrial-myelin associations ($R^2 = 0.025$-$0.309$; Fig. S15A). The variability of these regression slopes across conditions is further illustrated by bootstrap resampling (Fig. S22). These weak-to-moderate associations are compatible with geometric co-scaling effects, in which images containing a higher proportion of large-diameter axons tend to exhibit increased mitochondrial and myelin area fractions. However, image-level regressions alone do not resolve whether mitochondrial content and myelin area are directly related or whether their co-variation reflects broader genotype-dependent shifts in baseline ultrastructural composition. Nevertheless, the cross-genotype trend aligns with the broader pattern previously observed in (Fig. 4), where mitochondrial content tends to increase under conditions of reduced *Mbp* expression.
Axon-myelin regressions also displayed uniformly positive slopes across all genotypes (Fig. S15B). The strength of association varied substantially, with the triple KO showing a particularly high coefficient of determination ($R^2 = 0.811$), reflecting a more constrained structural regime, with relatively uniform populations of small, hypomyelinated axons. Other genotypes exhibited weaker correlations ($R^2 = 0.089$-$0.362$), likely reflecting increased heterogeneity in axonal caliber distributions and overall field composition.
The mitochondrial-myelin and axon-myelin regressions here suggest that their co-variation at the image level may reflect a combination of shared geometric scaling with axonal size and genotype-dependent shifts in baseline ultrastructural composition. Importantly, because these measurements are aggregated at the image level, mitochondrial area fractions include contributions from both myelinated and unmyelinated axons, whereas myelin area reflects only myelinated fibers, such that the observed associations are also influenced by overall tissue composition. These population-level patterns therefore reflect a combination of geometric scaling and tissue composition effects, rather than direct mechanistic coupling, motivating more targeted per-fiber analyses.
To further assess the stability and genotype dependence of image-level associations between mitochondrial and myelin area fractions, we estimated regression slopes using bootstrap resampling (Fig. S22).
Across genotypes, regression slopes remain positive but exhibit a non-monotonic pattern, with an increase in single KO conditions followed by a reduction in more severe KO groups. This behavior suggests that the apparent strength of mitochondria–myelin coupling varies across conditions and is not simply proportional to the degree of *Mbp* perturbation.
One possible explanation is that, in mild perturbations, shifts in tissue composition or global mitochondrial content may enhance the apparent correlation at the image level. In contrast, under more severe *Mbp* deficiency, increased structural heterogeneity and reduced myelin content may weaken this relationship by limiting the dynamic range of myelin-associated variation.
Interestingly, this genotype-dependent variability is similar to subsequent per-fiber and diameter-controlled analyses (Fig. S23), which show that mito-myelin associations are not stable at the level of individual axons and are strongly influenced by geometric and compositional factors.

***A.2. Morphometric feature extraction.*** For each segmented object instance, we extracted a set of label-specific morphometric descriptors defined in a curated lookup table. The metrics included in the analysis were selected based on their relevance to ultrastructural geometry, robustness to segmentation noise, and interpretability for downstream statistical modeling.

In order to have label-specific instance metrics, each semantic class was associated with a predefined set of shape and area descriptors:

**Axon, mitochondria, mitochondria outside axon, mito-like organelles:** *AbsoluteArea* ($\mu$m$^2$), *Perimeter*, *Solidity*, *Eccentricity*, *Circularity*, *EquivalentDiameter*.

**Myelin, periaxonal space (PAS), abnormality:** *AbsoluteArea* ($\mu$m$^2$), *Perimeter*, *Solidity*.

AbsoluteArea here denotes the object's physical cross-sectional area after conversion from pixels using the image-specific spatial calibration. Shape descriptors were derived from standard region properties.
To obtain image-level representations, instance-level measurements were aggregated using a fixed set of summary statistics:

**Central tendency:** mean, median

**Dispersion:** standard deviation

**Distribution shape:** skewness, kurtosis

**Percentiles:** 10th and 90th percentiles

**Extrema:** maximum

**Size descriptors:** instance count and total area

**Normality indicator:** binary flag based on Shapiro-wilk or D'Agostino test, depending on sample size

These aggregated statistics formed a comprehensive image-level feature set for each semantic label and were subsequently used for downstream analyses, including visualization and genotype prediction.

***A.3. Random forest for each ultrastructure.*** To explore the relative contribution of different ultrastructural compartments to genotype discrimination, we trained separate random forest classifiers using feature subsets derived from axons, myelin, mitochondria, and periaxonal space.

Among the compartment-specific models, myelin-derived features yielded the strongest classification performance overall (Fig. S16). Severe enhancer deletions remained well separated, and the dominant predictors were myelin area, perimeter, and solidity descriptors, in agreement with the central role of myelin remodeling in shaping genotype-specific ultrastructural phenotypes.

Axon-derived features also retained substantial discriminative information, but with broader overlap among intermediate and mild perturbations (Fig. S17). Their most informative variables were dominated by caliber, count, and shape related descriptors, shows that axonal geometry do contributes to genotype separation, although less strongly than myelin-specific features.

Periaxonal space-derived features showed intermediate-to-strong discriminative power, exceeding that of mitochondria-derived features and preserving clear separation of the most severe genotype (Fig. S19). This apparent separability likely reflects substantial morphological remodeling of the periaxonal space under severe *Mbp* deficiency. Rather than indicating a loss of PAS, these changes are consistent with disruption of its canonical "tongue-like" organization, which becomes replaced or obscured by unmyelinated profiles, membrane irregularities, and overlapping structural components. As a result, PAS-associated features capture large-scale structural alterations in severe genotypes, while classification performance for intermediate and mild conditions remains limited.

In contrast, mitochondria-derived features produced a weaker overall classifier, with certain extent overlap across genotypes (Fig. S18). Nevertheless, they retained meaningful structure, particularly in distinguishing broad severity levels. Classification of double and triple KO groups remained relatively stable, and predictions rarely spanned distant genotype classes (e.g., severe KO conditions being misclassified as WT or M3KOKI). This constrained misclassification pattern suggests that mitochondrial features encode a continuous gradient of ultrastructural change that tracks the extent of *Mbp* perturbation, even though their standalone discriminative power is limited.

Notably, across axon-, mitochondria-, and PAS-restricted models, intermediate phenotypes were repeatedly attracted toward the M3KOKI class, indicating that this group occupies a broad region of ultrastructural feature space even when classification is based on a single compartment. By comparison, the myelin-only model reduced, but did not entirely eliminate, this ambiguity.

Across all compartment-specific models, classification performance for the single KO group should be interpreted with caution, as multiple distinct genotypes (M3KO, M3(225)KO, M5KO) were merged into a single class.

***A.4. Individual level correlation.*** To characterize ultrastructural scaling relationships and to establish the basis for subsequent controlled analyses, we assessed the dependence of g-ratio (Fig. S20) and mitochondrial area (Fig. S21) on fiber diameter (log-transformed) across spinal cord regions (CST, GC, and VM), genotypes, and ages. All analyses were restricted to myelinated fibers.

Across all conditions, both g-ratio and mitochondrial area exhibit clear associations with fiber diameter, in line with geometric scaling effects. Larger fibers tend to be associated with increased mitochondrial content, while g-ratio shows a more modest and region-dependent relationship with fiber diameter, reflecting the complex interplay between axonal caliber and myelin thickness.

Focusing first on the g-ratio (Fig. S20), within myelinated fibers, progressive *Mbp* deletion is associated with an overall upward shift in g-ratio across genotypes, consistent with reduced myelin thickness. Age-related patterns exhibit greater variability across regions and genotypes; however, lower g-ratio values are frequently observed at P400, suggesting a late-stage trend toward myelin thickening. While this trend varies across certain conditions, the results indicate that myelin remodeling under *Mbp* perturbation is both heterogeneous and context-dependent.

For mitochondrial area (Fig. S21), within myelinated fibers, a positive association with fiber diameter is consistently observed, although with substantially greater dispersion compared to g-ratio. While WT and milder perturbation groups exhibit relatively compact scaling relationships, more severe KO conditions display increased variability in mitochondrial area among fibers of similar diameter. This indicates that mitochondrial content is influenced not only by geometric scaling but also by additional sources of variability under conditions of *Mbp* deficiency.

Together, these results identify fiber diameter as a major geometric determinant influencing both g-ratio and mitochondrial content. Crucially, because both variables co-vary with fiber size, simple correlations between mitochondrial measurements and g-ratio may largely reflect shared dependence on diameter rather than direct coupling. This motivates the use of diameter-controlled analyses, such as partial correlation, to isolate their intrinsic relationship (Fig. 10). The instability of the mitochondria-g-ratio relationship across conditions is further supported by bootstrapped slope estimates, which show substantial variability across regions, ages, and genotypes (Fig. S23).

***A.5. Bootstrapping mitochondria–g-ratio slope on fibers.*** To further assess the stability of the relationship between mitochondrial content and g-ratio within myelinated fibers, we estimated regression slopes using bootstrap resampling across regions, ages, and genotypes (Fig. S23).
Across all conditions, slope estimates exhibit substantial variability, with both magnitude and direction differing across regions and genotypes. In particular, slopes frequently approach zero or change sign across conditions, indicating that the association between mitochondrial area and g-ratio is not stable at the per-fiber level.
This lack of consistency suggests that apparent associations between mitochondrial content and g-ratio are unlikely to reflect a direct or universal coupling, but instead arise from context-dependent factors, including geometric scaling and population heterogeneity.

***A.6. Ultrastructural morphology in age progress.*** To characterize developmental trends in ultrastructural composition, we examined age-dependent trajectories of mitochondrial and myelin area fractions in the CST region (Fig. S24, Fig. S25).
Across genotypes, mitochondrial area fraction exhibits a non-monotonic trajectory in KO lines, with an increase from P30 to P90 followed by a decline toward P400 in conditions where later time points are available (double KO group), suggesting that mitochondrial expansion is temporally regulated and may be most pronounced at intermediate developmental stages. In contrast, WT and M3KOKI groups display a more stable or gradually decreasing trend, a more distinct developmental dynamics under preserved or partially restored *Mbp* expression.
By comparison, myelin area fraction shows a consistent increase across age in nearly all genotypes, reflecting ongoing myelination during development. Although severe KO groups remain shifted toward lower overall myelin levels, the upward trajectory is preserved, suggesting that developmental progression of myelin accumulation continues despite reduced *Mbp* expression.
These temporal trends provide additional context for the population-level and per-fiber analyses presented in the main text. Importantly, while age-dependent trends are observable, these effects are secondary to genotype-dependent shifts and do not alter the primary conclusions derived from age-pooled analyses in the main text.

***A.7. Heterogeneous mitochondrial presence across axons.*** Building on the observation that absolute mitochondrial area increases at the per-fiber level under severe KO conditions (Fig. 9A, B), we next asked whether the broader increase in mitochondrial content observed at the image level reflects a higher proportion of mitochondria-containing axons or greater mitochondrial accumulation within a specific subset of fibers (Fig. S26).
Across most genotypes, the fraction of axons containing at least one mitochondria shows only moderate variation. In the most severe KO conditions, this fraction exhibits a slight reduction, indicating that fewer individual axons contain detectable mitochondria. When considered together with the increase in per-fiber mitochondrial area, this pattern suggests that mitochondrial remodeling does not reflect a uniform expansion across the axonal population. Instead, the observed increase appears to be driven by greater mitochondrial accumulation within a subset of fibers, rather than by a broad increase in mitochondria-containing axons.

***A.8. Cross-individual evaluation.*** To assess whether the learned morphological representations reflect genotype-level characteristics rather than animal-specific features, we performed a leave-one-animal-out evaluation (Fig. S27). In this setting, all images from selected animals were excluded from training and validation and reserved exclusively for testing. Genotype-, age-, and region-matched conditions were retained in the training set to ensure comparable biological contexts.
The held-out dataset included three spinal cord regions (CST, GC, and VM) collected at P90 from the M3(3x43bp) line (genotypically equivalent to M3KOKI), as well as an independent experimental replicate of the M3KO P30 VM condition. These samples originated from distinct animals and therefore provided a direct test of cross-individual generalization.
Because not all held-out subclasses were represented in the training taxonomy, classification performance was summarized using a rectangular confusion matrix, with rows corresponding to test-set labels and columns representing trained genotype classes. Row normalization expresses each held-out condition as a probability distribution over predicted genotypes.
Predictions for held-out animals remained predominantly assigned to the correct genotype (Fig. S27), with misclassifications largely confined to closely related M3-derived lines. These results were obtained using the Swin-Large architecture, which is also employed as the encoder backbone in our segmentation framework. Together, this analysis supports the interpretation that the extracted morphological features capture stable genotype-dependent patterns that generalize across animals and experimental repeats.

#### B. Supplementary Figures.

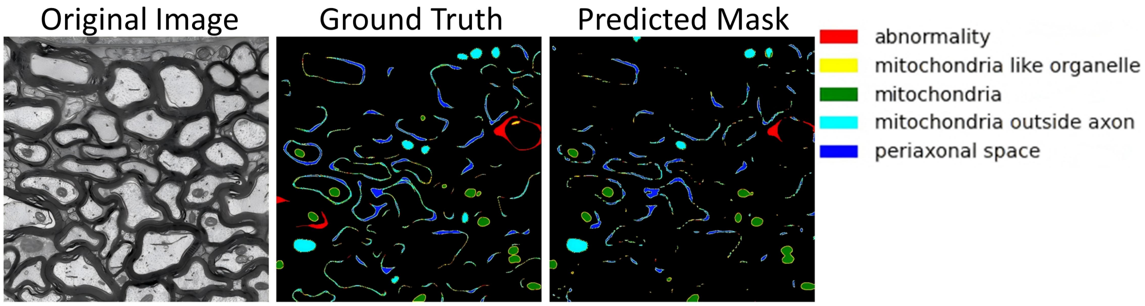


**Fig. S1.** Example of EM image segmentation comparing the ground truth annotation and the model prediction. Left: Original transmission electron microscopy (TEM) image. Middle: Ground-truth annotation with labeled classes, including abnormality (red), mitochondria-like organelles (yellow), mitochondria (green), mitochondria outside the axon (cyan), and periaxonal space (blue). Right: Predicted segmentation mask generated by the model.

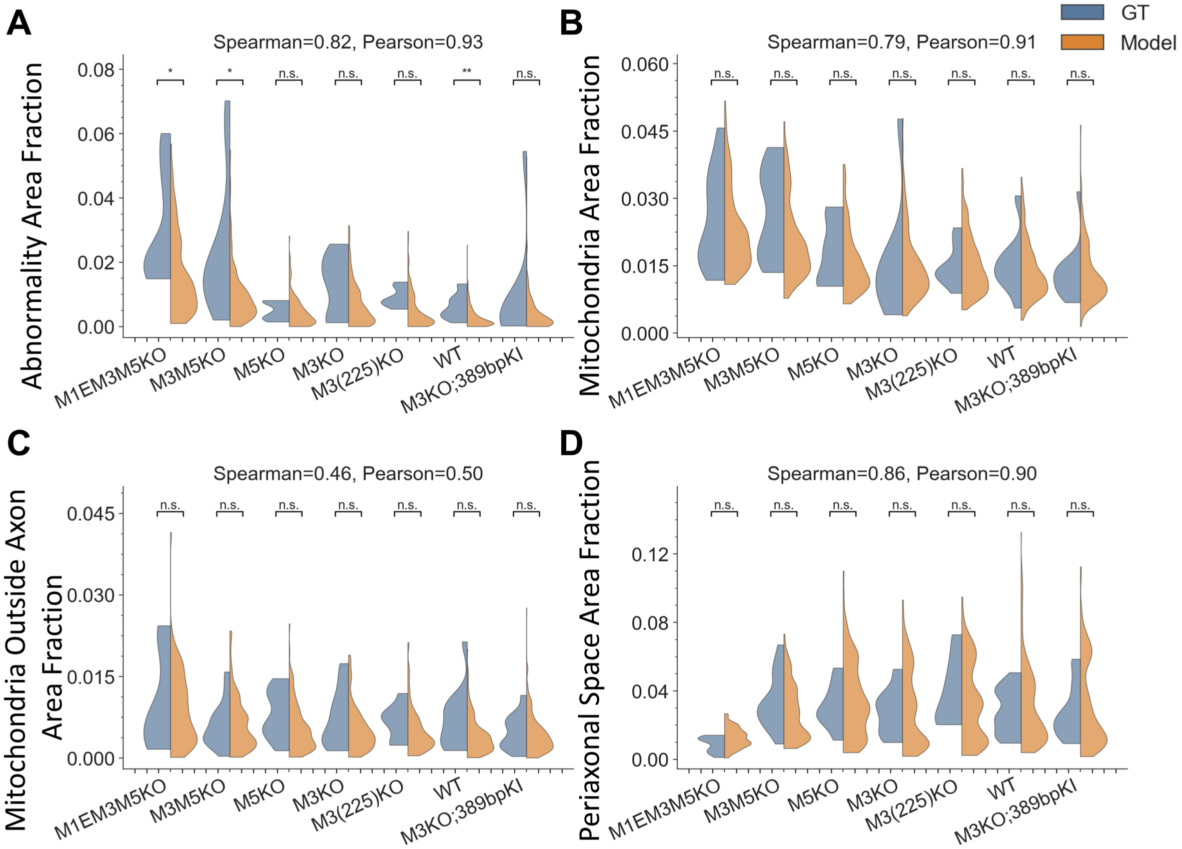


**Fig. S2. Comparison between ground truth and model predictions across ultrastructural metrics.** Violin plots of per-image distributions comparing ground truth (GT) annotations and model predictions across all genotypes and regions. **(A)** Area fraction comparison for abnormality. **(B)** Area fraction comparison for mitochondria. **(C)** Area fraction comparison for mitochondria outside of axon. **(D)** Area fraction comparison for periaxonal space. For each structure, split violins represent GT (blue) and model (orange) distributions, and significance was assessed using two-sided Mann-Whitney U tests with Holm-Bonferroni correction. Spearman and Pearson correlation coefficients report the agreement between GT and model medians across structures.

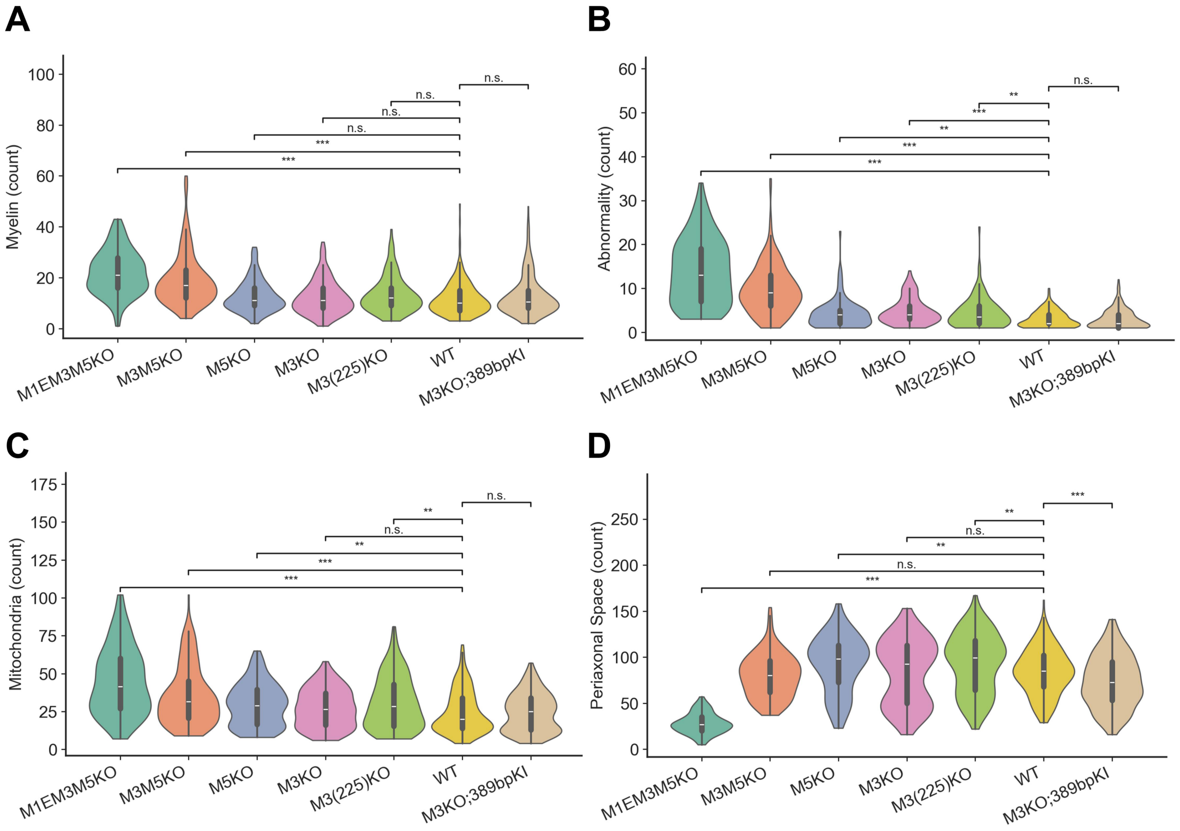


**Fig. S3. Pooled per-image distributions of ultrastructural counts across *Mbp* genotypes.** Violin plots of per-image distributions of ultrastructural counts of **(A)** myelin, **(B)** abnormality, **(C)** mitochondria and **(D)** periaxonal space.

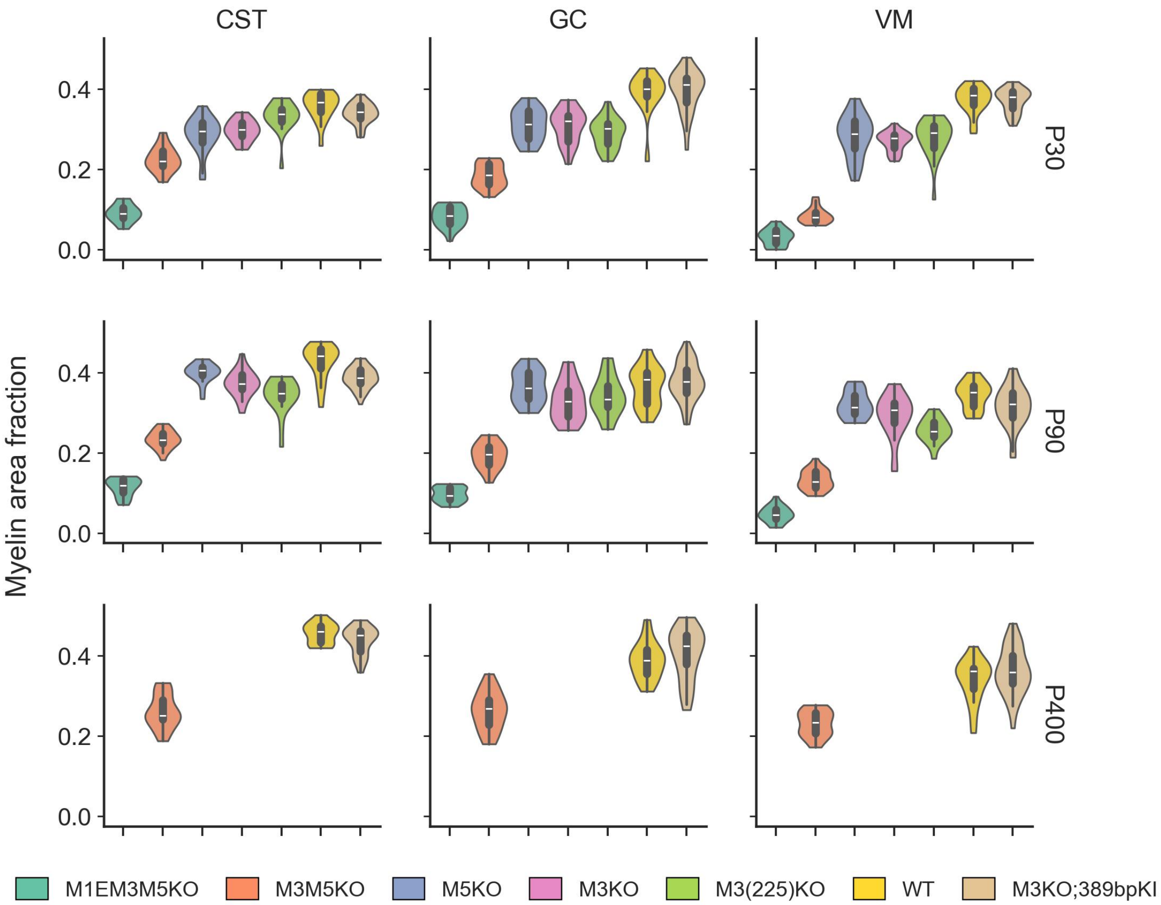


**Fig. S4. Distribution of myelin area fraction across *Mbp* genotypes, ages and spinal cord regions.** Violin plots show the total area per image ($\mu m^2$) for each genotype, faceted by age (rows: P30, P90, P400) and region (columns: CST, GC, VM). Each violin represents the distribution of measurements from each individual images, with embedded box plots indicating median and quartiles. No images were acquired at P400 for the single and triple KO conditions.

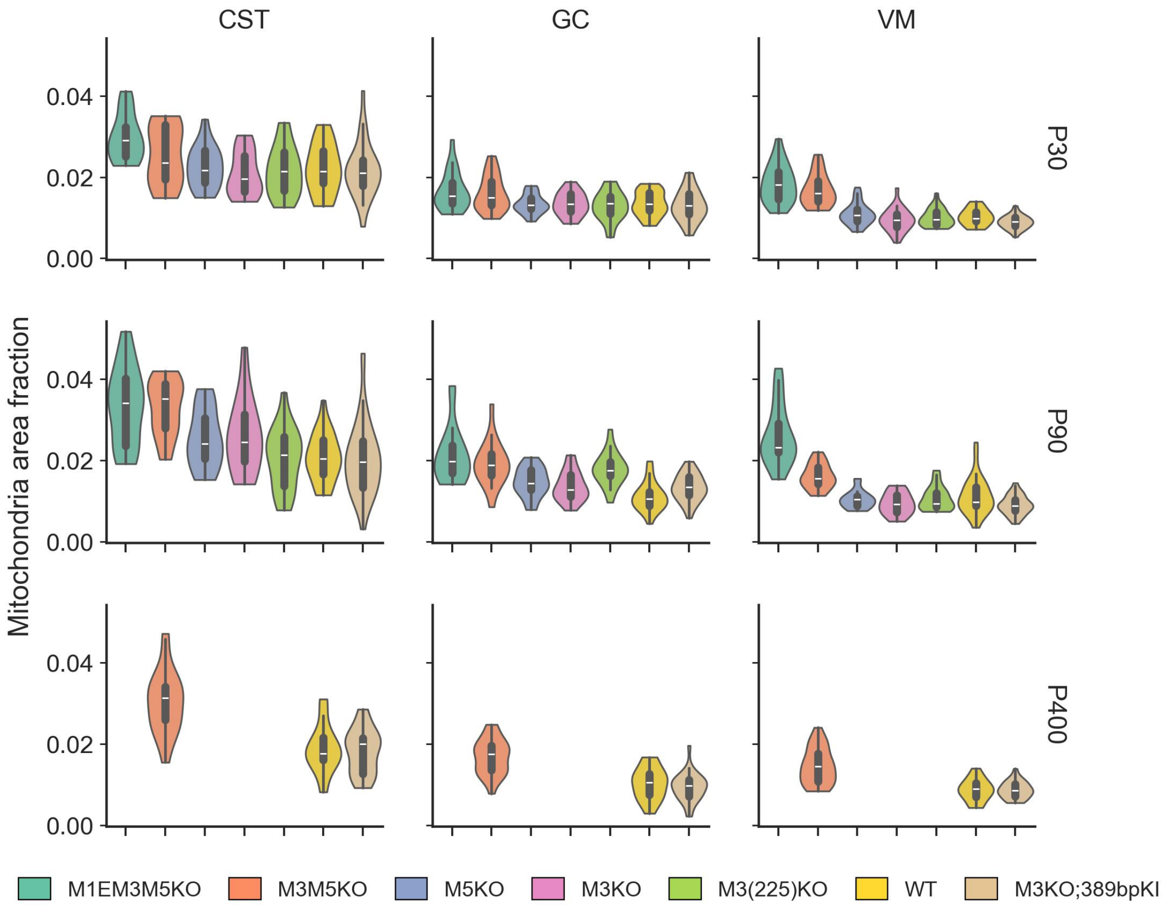


**Fig. S5. Distribution of mitochondrial area fraction across *Mbp* genotypes, ages, and spinal cord regions.** Violin plots show the total area per image ($\mu m^2$) for each genotype, faceted by age (rows: P30, P90, P400) and region (columns: CST, GC, VM). Each violin represents the distribution of measurements from individual images, with embedded box plots indicating median and quartiles. No images were acquired at P400 for the single and triple KO conditions.

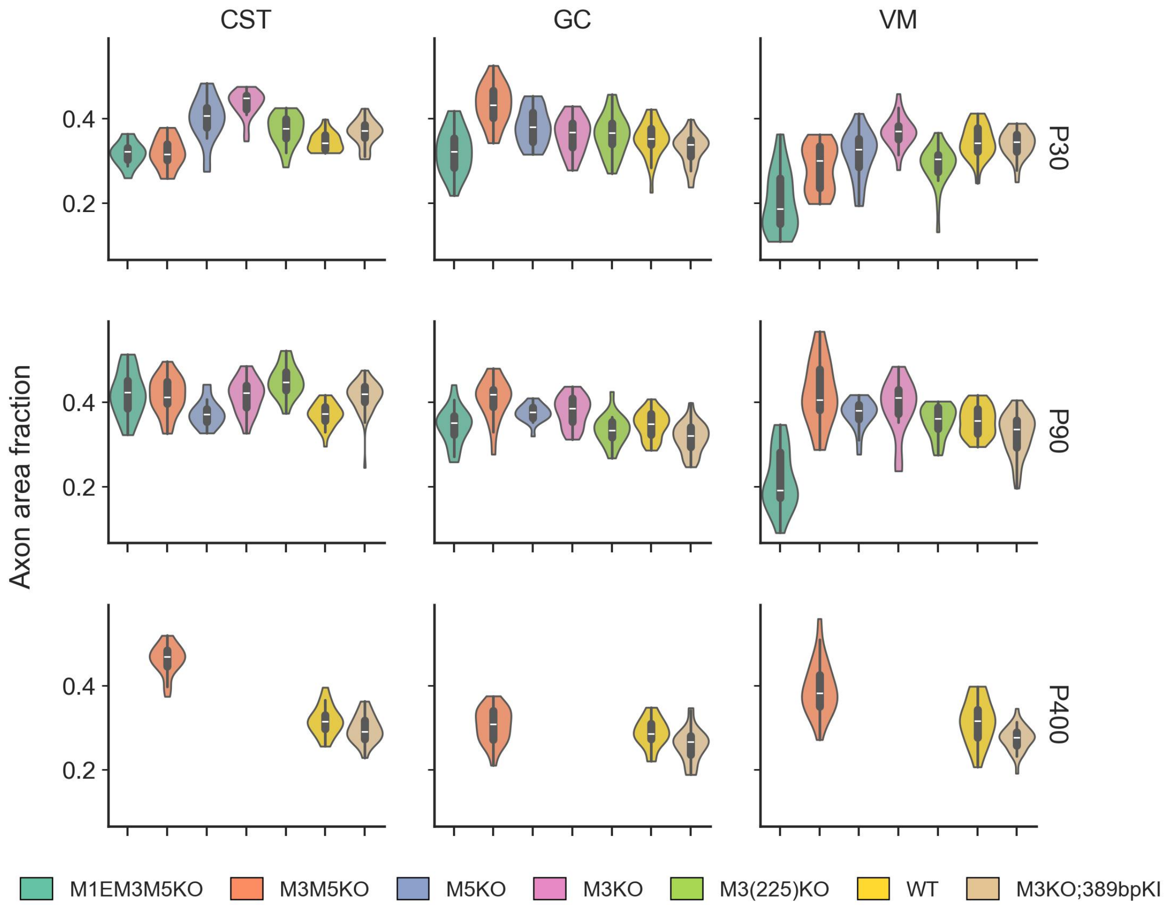


**Fig. S6. Distribution of axon area fraction across *Mbp* genotypes, ages, and spinal cord regions.** Violin plots show the total area per image ($\mu m^2$) for each genotype, faceted by age (rows: P30, P90, P400) and region (columns: CST, GC, VM). Each violin represents the distribution of measurements from individual images, with embedded box plots indicating median and quartiles. No images were acquired at P400 for the single and triple KO conditions.

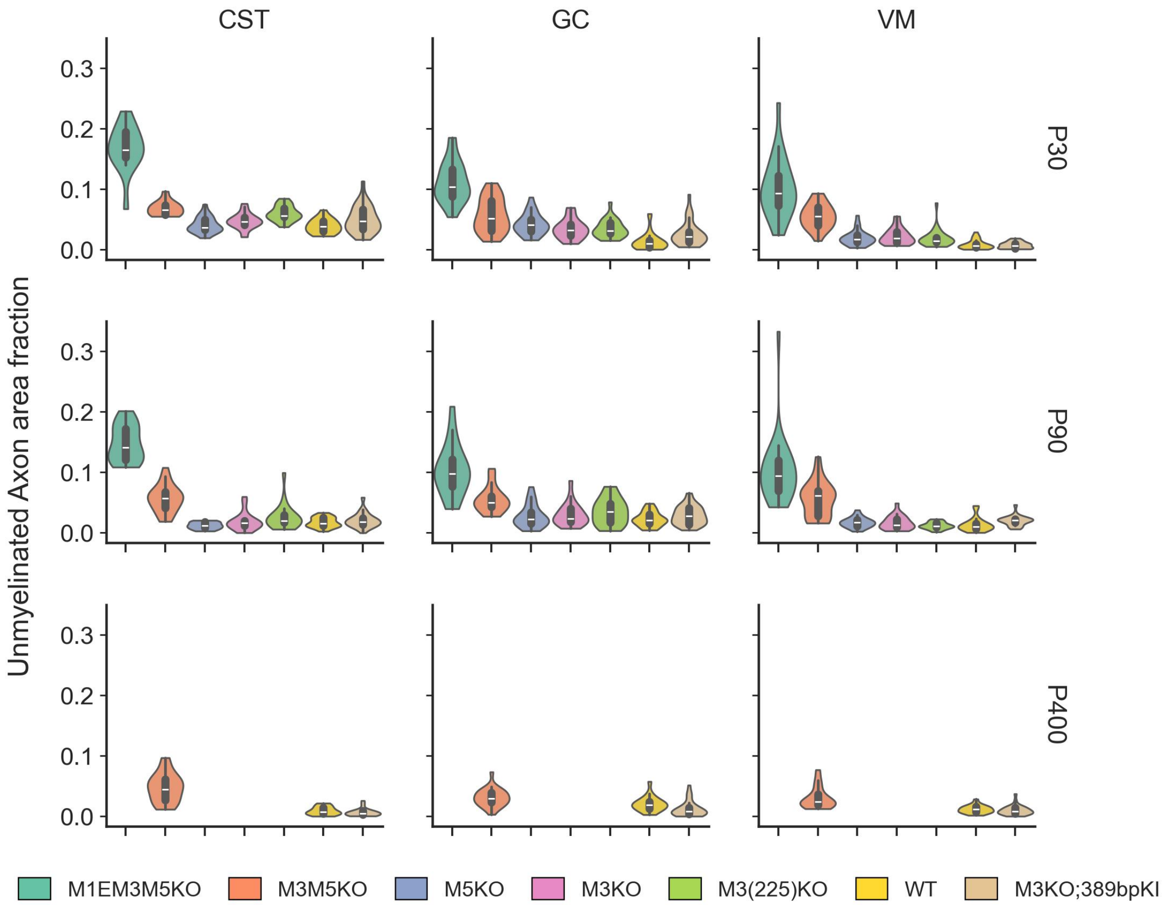


**Fig. S7. Distribution of unmyelinated axon area fraction across *Mbp* genotypes, ages, and spinal cord regions.** Violin plots show the total area per image ($\mu m^2$) for each genotype, faceted by age (rows: P30, P90, P400) and region (columns: CST, GC, VM). Each violin represents the distribution of measurements from individual images, with embedded box plots indicating median and quartiles. No images were acquired at P400 for the single and triple KO conditions.

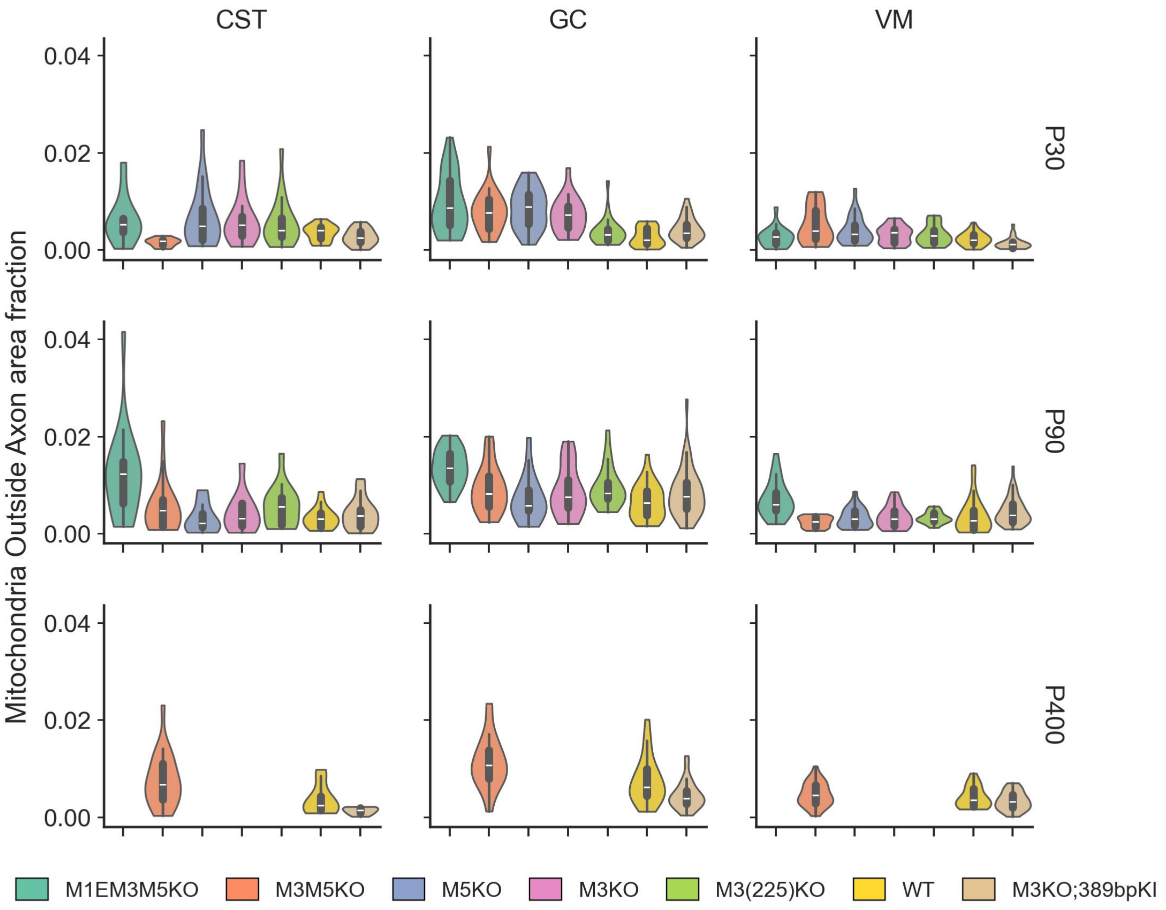


**Fig. S8. Distribution of mitochondrial area fraction outside axons across *Mbp* genotypes, ages, and spinal cord regions.** Violin plots show the total area per image ($\mu$m$^2$) for each genotype, faceted by age (rows: P30, P90, P400) and region (columns: CST, GC, VM). Each violin represents the distribution of measurements from individual images, with embedded box plots indicating median and quartiles. No images were acquired at P400 for the single and triple KO conditions.

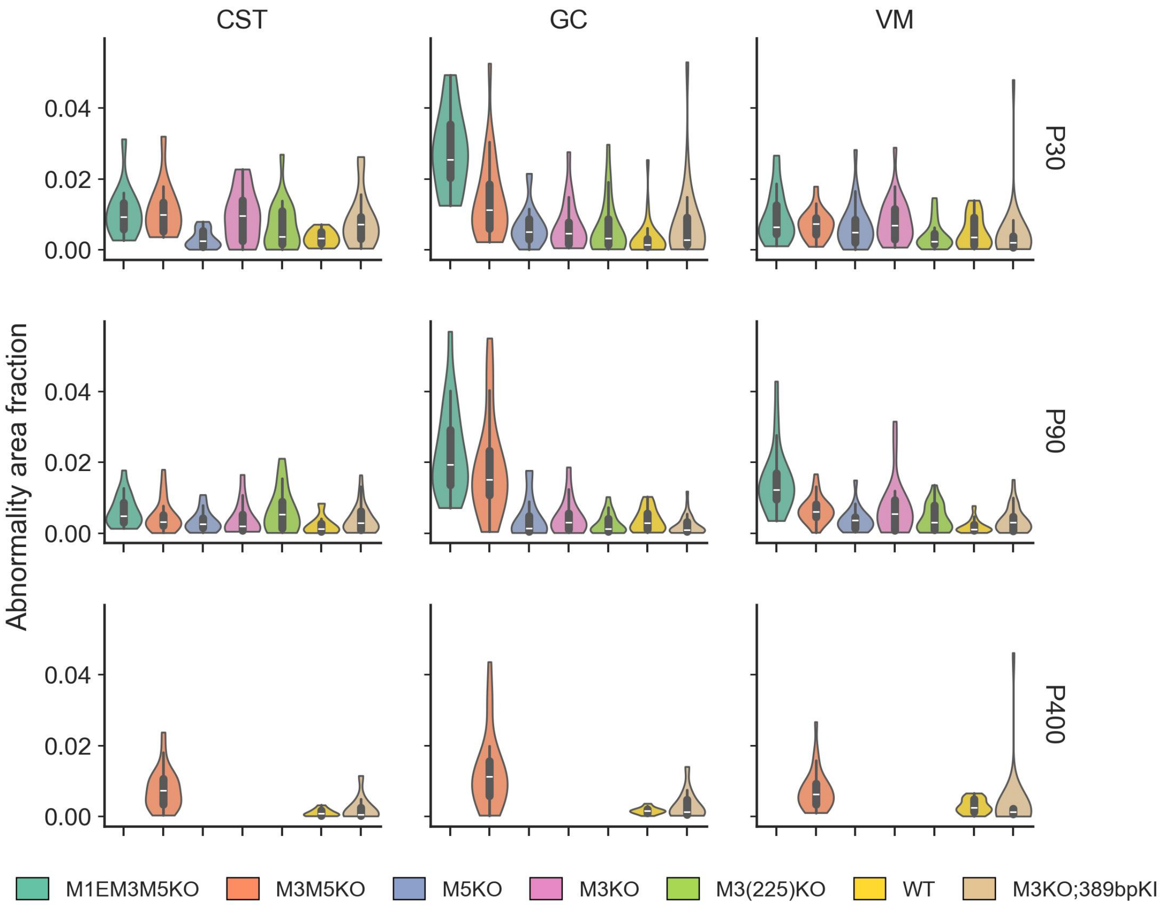


**Fig. S9. Distribution of abnormality area fraction across *Mbp* genotypes, ages, and spinal cord regions.** Violin plots show the total area per image ($\mu m^2$) for each genotype, faceted by age (rows: P30, P90, P400) and region (columns: CST, GC, VM). Each violin represents the distribution of measurements from individual images, with embedded box plots indicating median and quartiles. No images were acquired at P400 for the single and triple KO conditions.

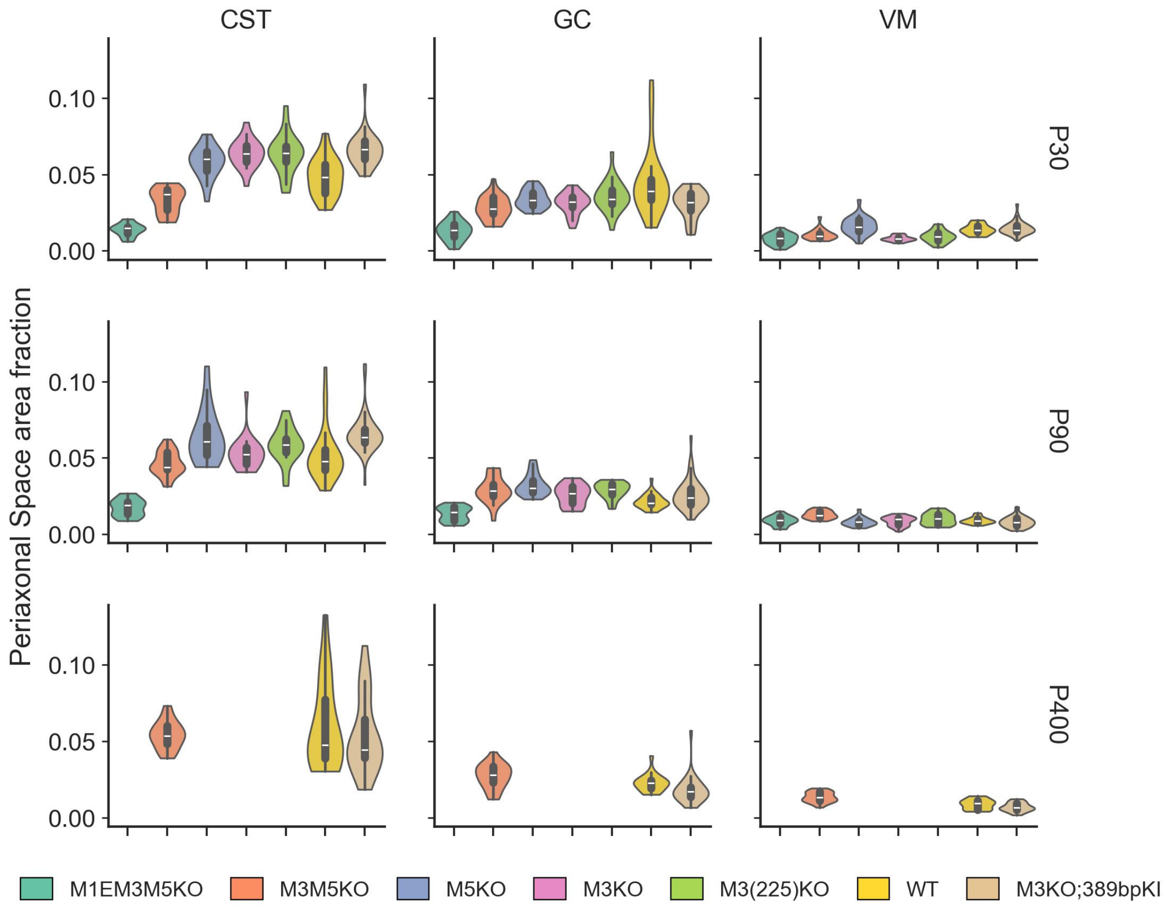


**Fig. S10. Distribution of periaxonal space area fraction across *Mbp* genotypes, ages, and spinal cord regions.** Violin plots show the total area per image ($\mu$m$^2$) for each genotype, faceted by age (rows: P30, P90, P400) and region (columns: CST, GC, VM). Each violin represents the distribution of measurements from individual images, with embedded box plots indicating median and quartiles. No images were acquired at P400 for the single and triple KO conditions.

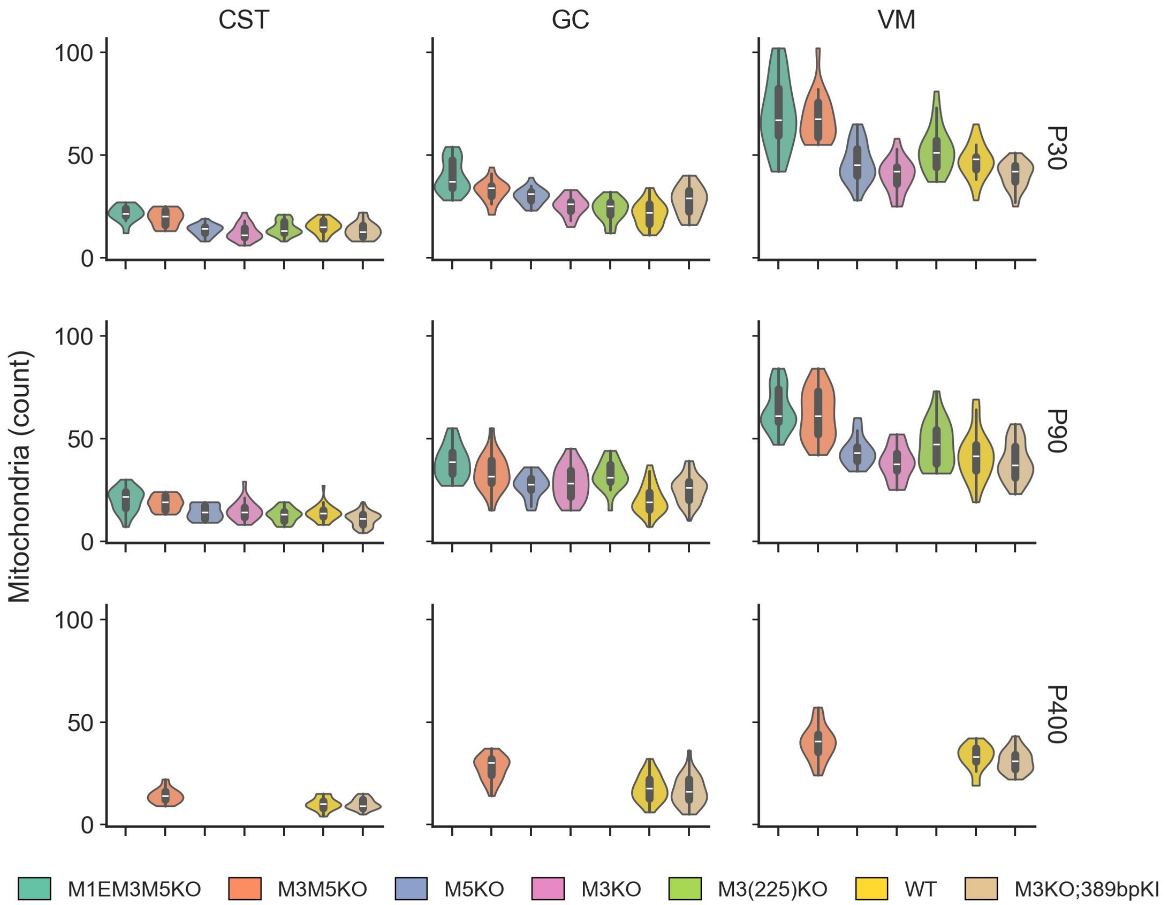


**Fig. S11. Distribution of mitochondria count across *Mbp* genotypes, ages, and spinal cord regions.** Violin plots show the total instance count per image for each genotype, faceted by age (rows: P30, P90, P400) and region (columns: CST, GC, VM). Each violin represents the distribution of measurements from individual images, with embedded box plots indicating median and quartiles. No images were acquired at P400 for the single and triple KO conditions.

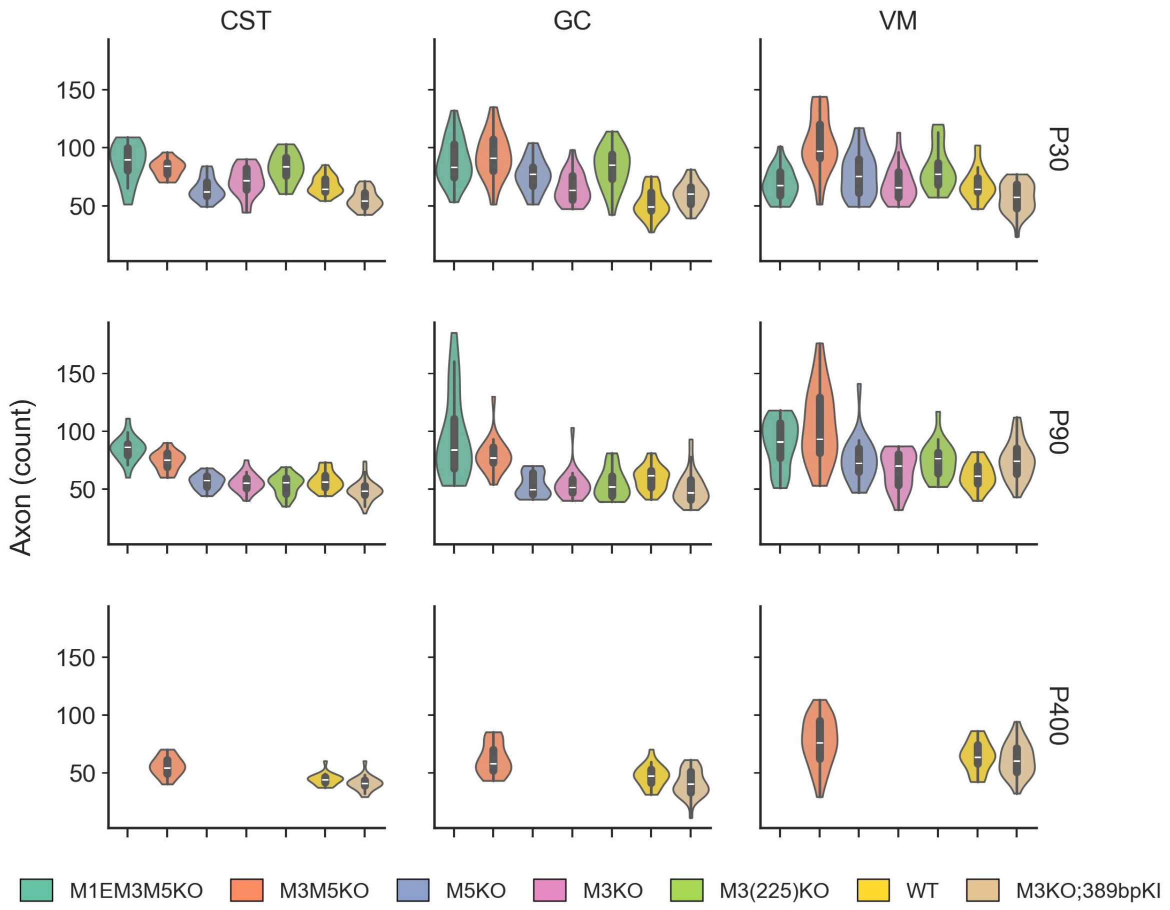


**Fig. S12. Distribution of axon count across *Mbp* genotypes, ages, and spinal cord regions.** Violin plots show the total instance count per image for each genotype, faceted by age (rows: P30, P90, P400) and region (columns: CST, GC, VM). Each violin represents the distribution of measurements from individual images, with embedded box plots indicating median and quartiles. No images were acquired at P400 for the single and triple KO conditions.

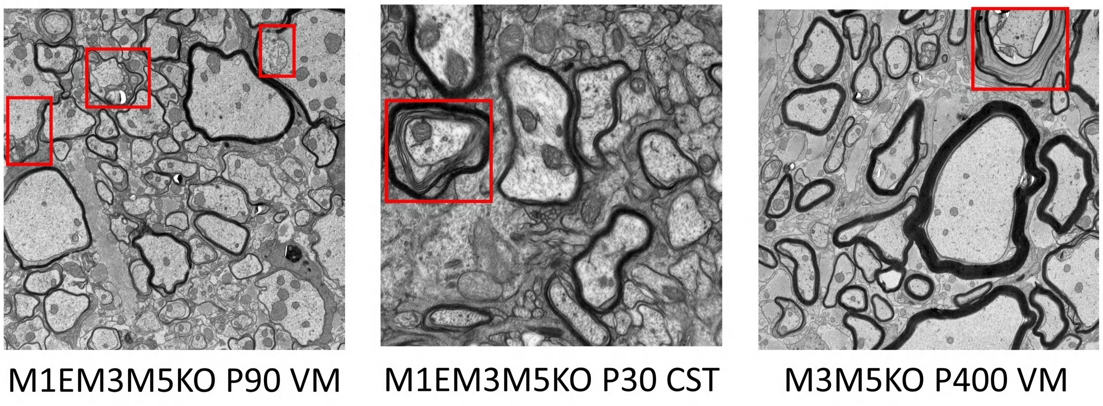


**Fig. S13. Examples of fibers exhibiting atypical morphology.** Representative examples displaying visually identifiable deviations from typical axon, myelin and PAS morphology. These examples are intended to illustrate qualitatively distinct structural patterns observed in the dataset. Highlighted regions indicate selected representative features, but do not exhaust the range of atypical morphologies present, which may also appear in other forms throughout the images.

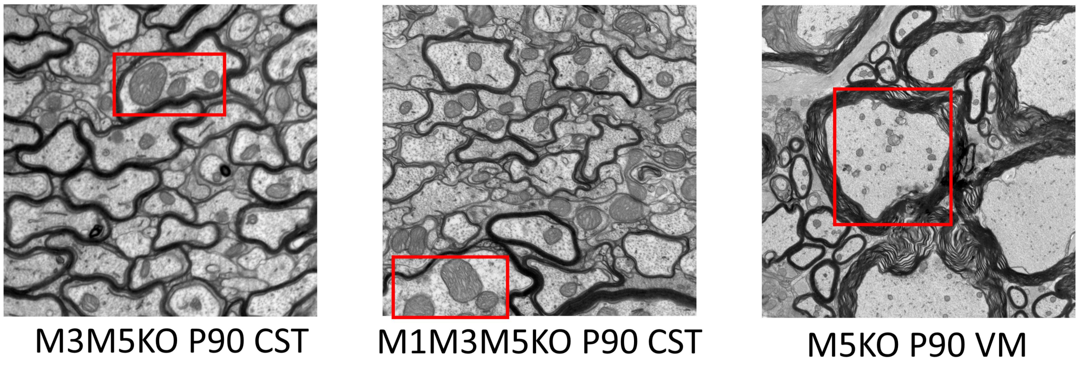


**Fig. S14. Fibers with elevated mitochondrial content.** Examples of fibers exhibiting unusually high mitochondrial area fractions or mitochondrial counts, identified based on statistical criteria.

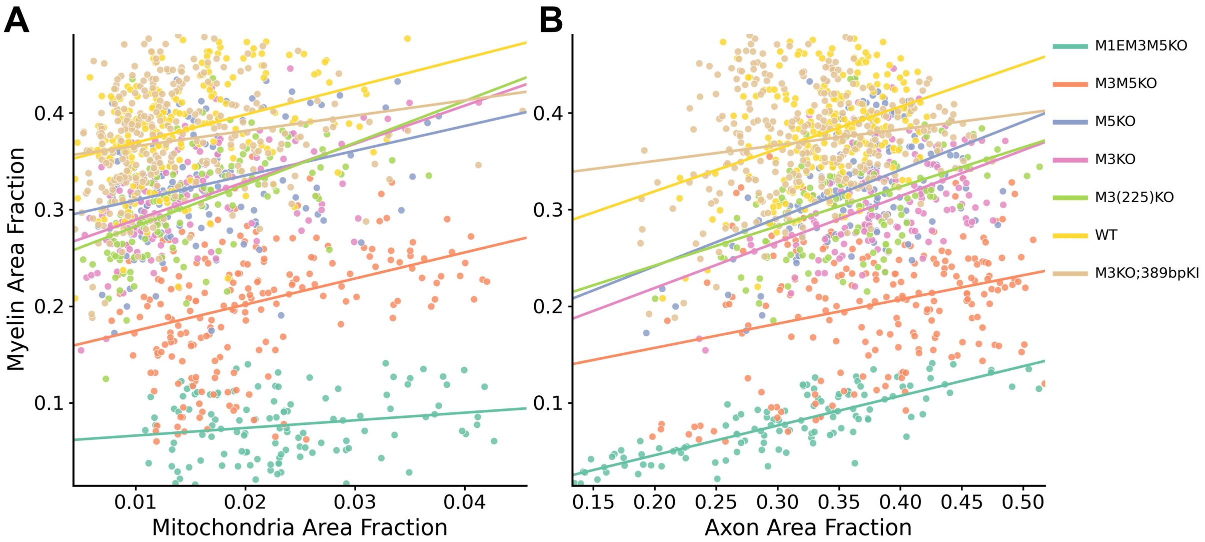


**Fig. S15. Cross-genotype correlations between mitochondrial and myelin area fractions pooled across ages and regions (per image). (A)** Mitochondrial versus myelin area fractions. **(B)** Axon versus myelin area fractions. For every genotype in each panel, linear regression is fitted using non-training (hold-out) images only, while hollow circles indicate ground-truth training images. Notice how across genotypes, both mitochondrial and axonal area fractions exhibit positive associations with myelin area fraction, with substantially stronger and more consistent correlations observed for axons (up to $R^2 = 0.811$). Full regression parameters for all genotypes are provided in Table 1 and Table 2.

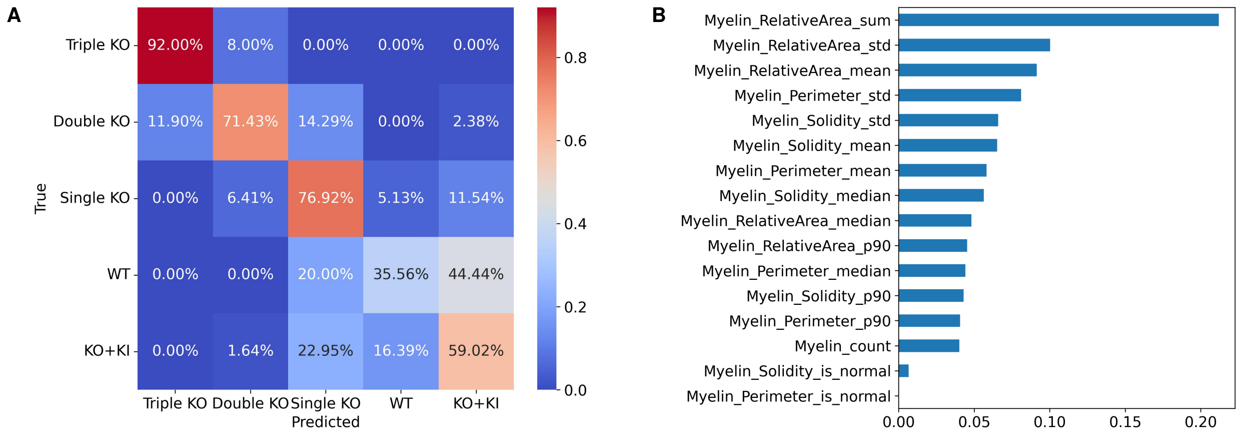


**Fig. S16. Random forest classification using myelin-derived features in a pooled analysis across ages and regions. (A)** Confusion matrix showing improved genotype separability compared to axon-only features, particularly for severe enhancer deletions, while overlap persists among milder perturbations. **(B)** Feature importance ranking highlighting myelin-related metrics, including relative area, perimeter, and solidity, as dominant predictors of genotype.

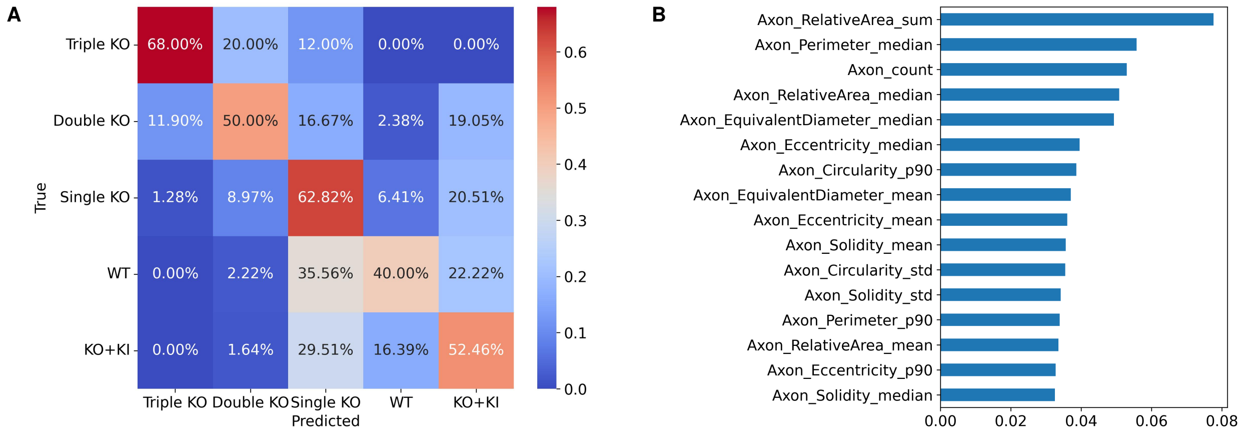


**Fig. S17. Random forest classification using axon-derived features in a pooled analysis across ages and regions. (A)** Confusion matrix showing genotype classification based solely on axonal morphometrics. Severe genotypes remain partially separable, but increased overlap is observed among intermediate and mild perturbations, with frequent misclassification toward the M3KOKI group. **(B)** Top features ranked by mean decrease in impurity (MDI), dominated by axon caliber and shape descriptors (e.g., relative area, diameter, circularity, and eccentricity).

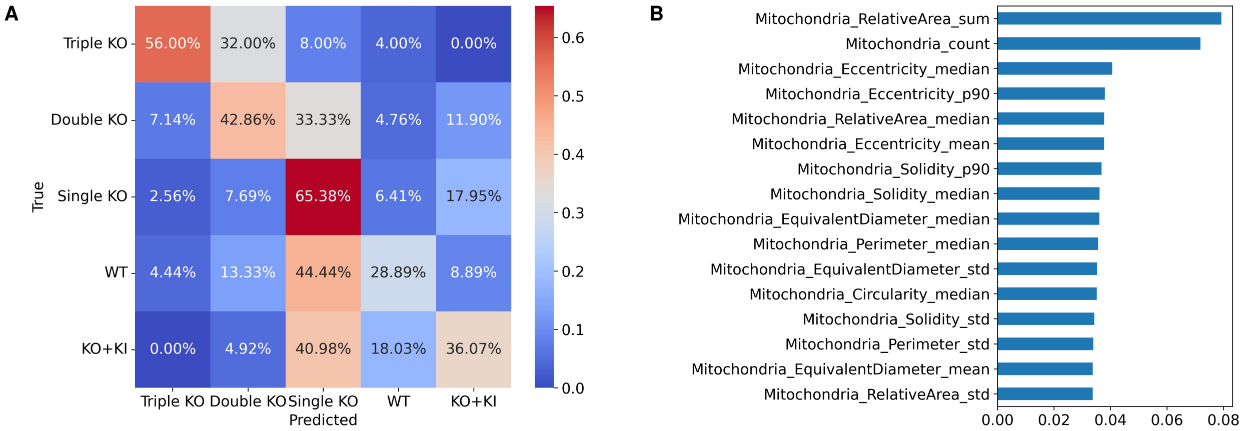


**Fig. S18. Random forest classification using mitochondrial features in a pooled analysis across ages and regions. (A)** Confusion matrix showing limited genotype separability, with substantial overlap across groups and a tendency for intermediate phenotypes to be assigned to the M3KOKI class. **(B)** Feature importance ranking indicating contributions from mitochondrial size, shape, and distribution descriptors, though with weaker discriminative power compared to myelin features.

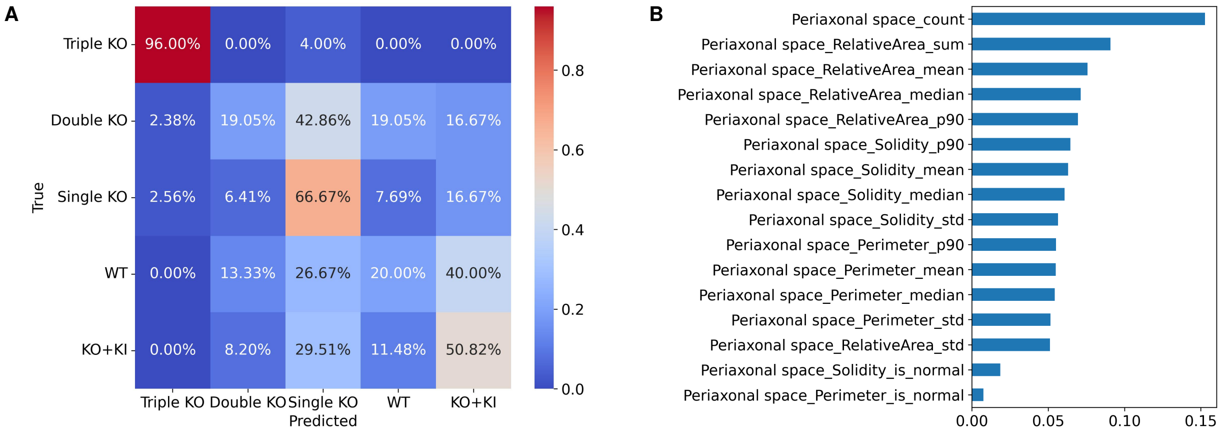


**Fig. S19. Random forest classification using periaxonal space features in a pooled analysis across ages and regions. (A)** Confusion matrix showing moderate classification performance, with partial separation of severe genotypes and increased overlap among milder conditions. **(B)** Feature importance ranking highlighting PAS-related metrics, including area, count, and relative occupancy, as contributing but secondary predictors.

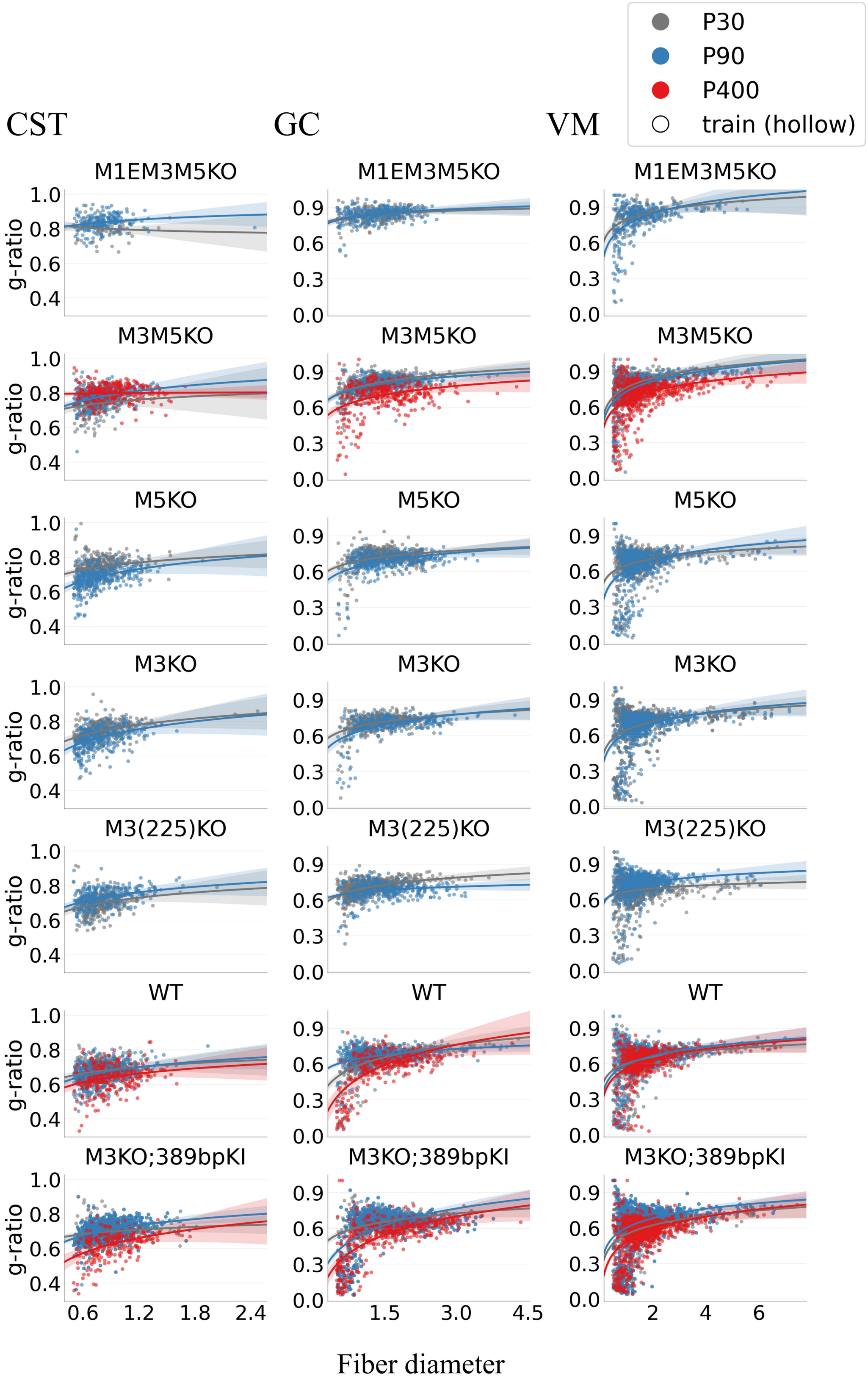


**Fig. S20. Relationship between fiber diameter ($\mu$ m) and g-ratio across *Mbp* genotypes, ages, and spinal cord regions.** Each panel shows per-fiber measurements with linear regression fits. Unmyelinated fibers are excluded. Across conditions, g-ratio exhibits an approximately linear dependence on log-transformed fiber diameter, match with geometric scaling effects. Detailed regression statistics for each age, genotype, and region are provided in Table 3.

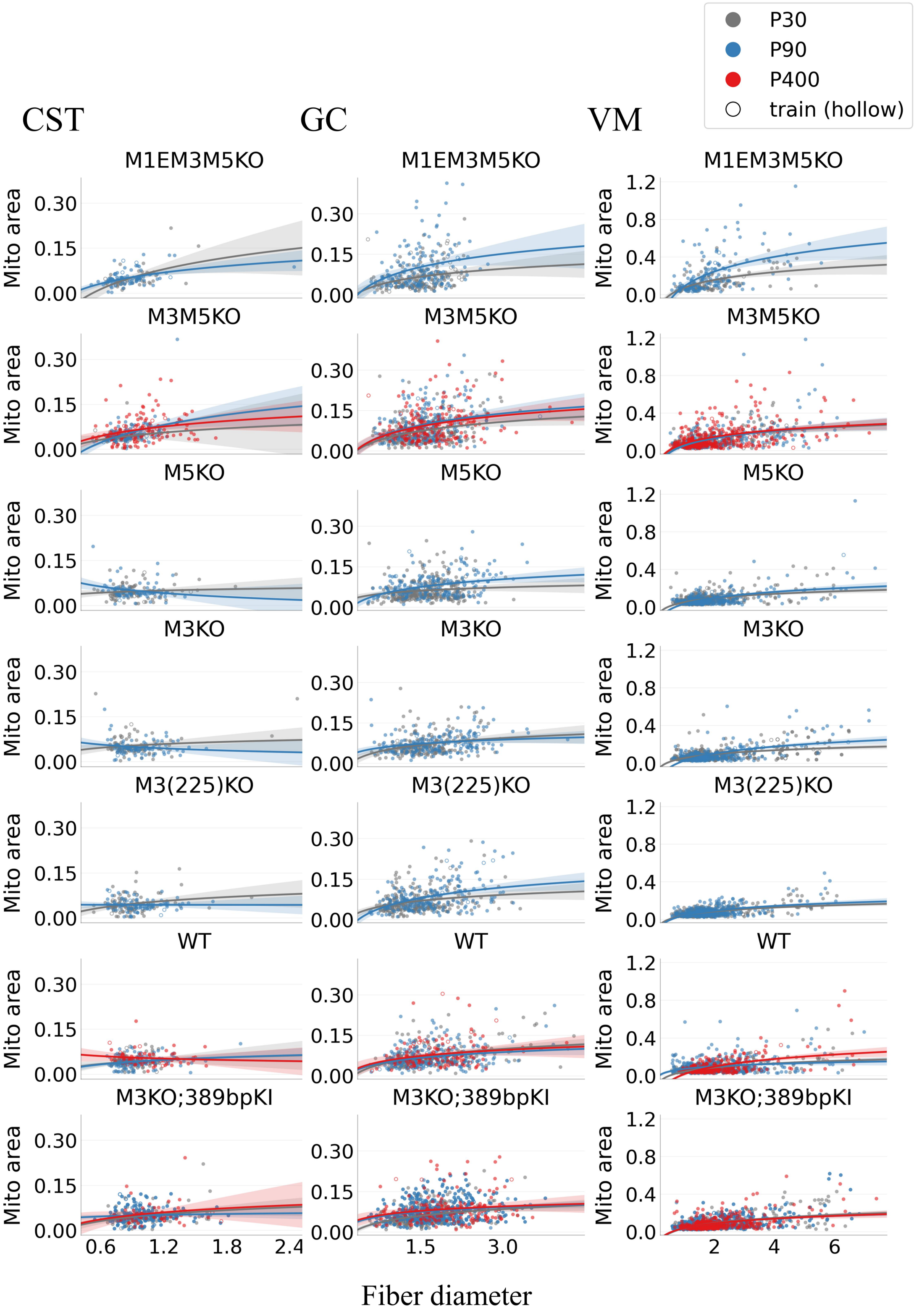


**Fig. S21. Relationship between fiber diameter ($\mu m$) and mitochondrial area ($\mu m^2$) across *Mbp* genotypes, ages, and spinal cord regions.** Each panel shows per-fiber measurements with linear regression fits. Unmyelinated fibers are excluded. Across conditions, mitochondrial area exhibits an approximately linear dependence on log-transformed fiber diameter, match with geometric scaling effects. Detailed regression statistics for each age, genotype, and region are provided in Table 4.

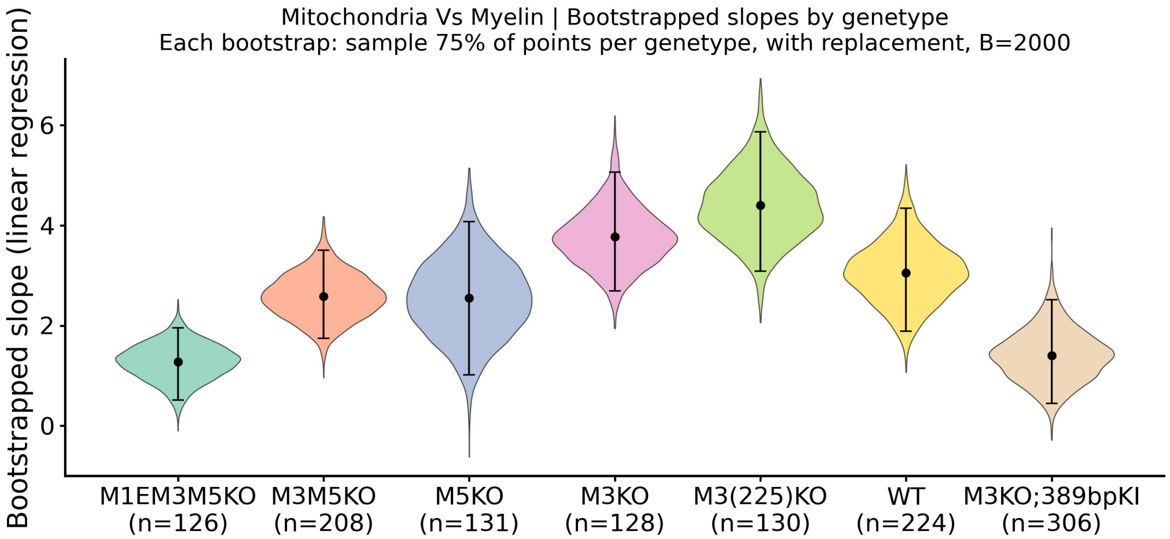


**Fig. S22. Bootstrap estimates of image-level mitochondria-myelin regression slopes across *Mbp* genotypes, pooled across ages and regions.** Linear regression slopes between mitochondrial and myelin area fractions were estimated using bootstrap resampling with 2000 bootstrap replicates ($B = 2000$). While slopes are consistently positive, their magnitudes exhibit a non-monotonic pattern across genotypes, increasing in single KO conditions and decreasing in more severe KO groups. The spread of the distributions further indicates substantial variability in slope estimates across conditions.

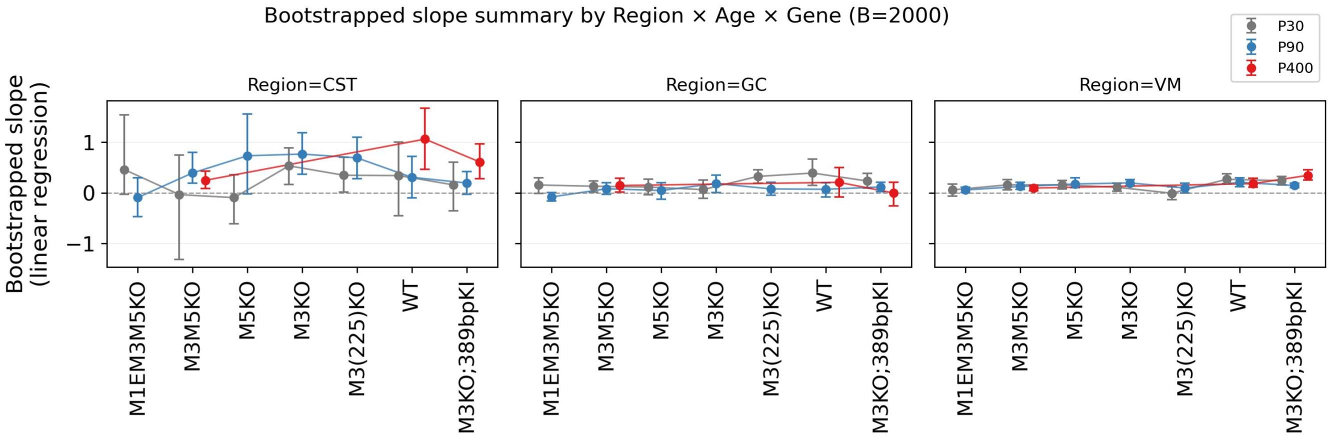


**Fig. S23. Bootstrap estimates of the mitochondia–g-ratio regression slope across genotypes, ages and regions.** Linear regression slopes between mitochondrial area and g-ratio were estimated using bootstrap resampling with 2000 bootstrap replicates ($B = 2000$) across regions, ages, and *Mbp* genotypes. Error bars indicate confidence intervals. Analysis restricted to myelinated fibers. Slopes exhibit substantial variability across conditions, with no consistent trend across genotypes or regions, indicating that the mitochondria–g-ratio relationship is not stable and may be strongly influenced by underlying heterogeneity.

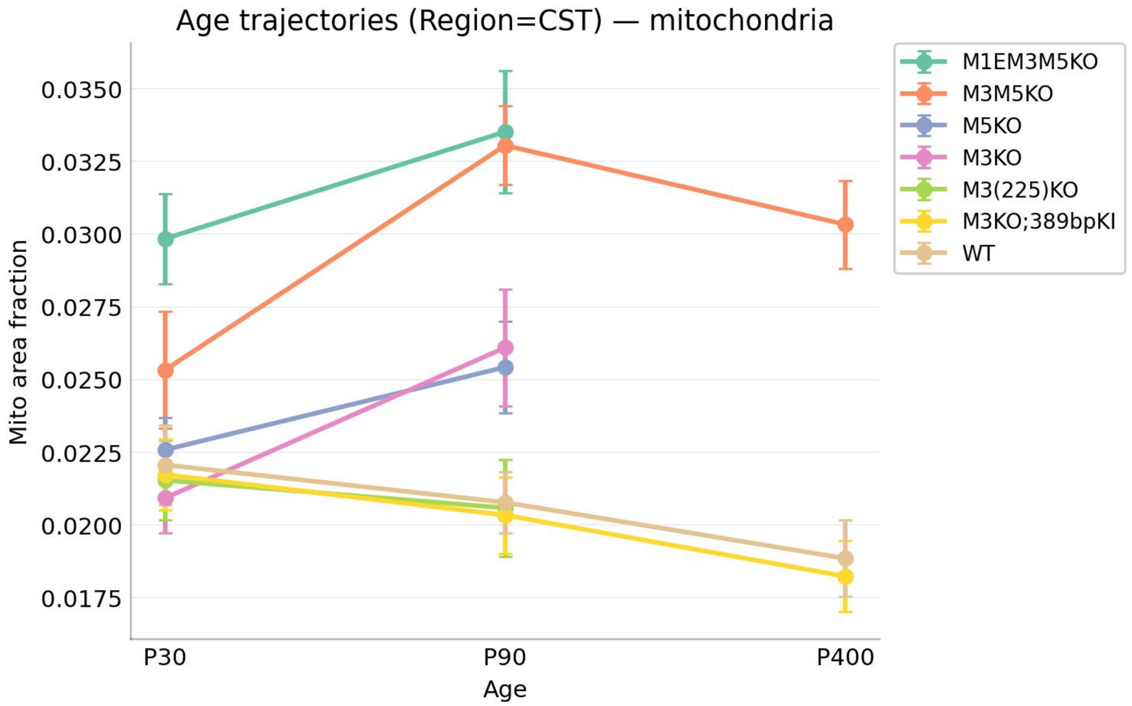


**Fig. S24. Age trajectories of mitochondrial area fraction across *Mbp* genotypes in the CST region.** Data represent the mean mitochondrial area fraction per image, aggregated across all analyzed sections. Results are presented as $Mean \pm SEM$ for each genotype at P30, P90, and P400. KO lines exhibit an initial increase in mitochondrial area between P30 and P90, followed by a decline by P400 (available for double KO only). In contrast, the WT and M3KOKI groups display a distinct developmental pattern.

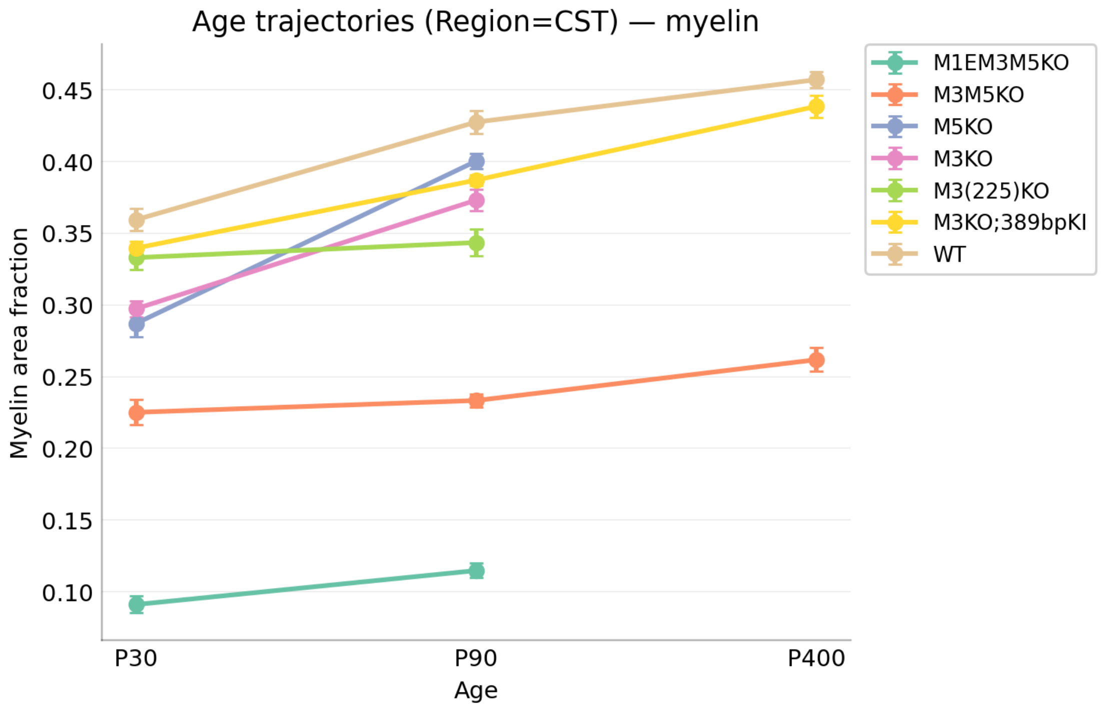


**Fig. S25. Age trajectories of myelin area fraction across *Mbp* genotypes in the CST region.** Data represent the mean area fraction per image, aggregated across all analyzed sections. Values are expressed as $Mean \pm SEM$ for each genotype at P30, P90, and P400. Nearly all mouse lines exhibit a progressive increase in myelin area fraction throughout development, suggesting that the myelination process proceeds regardless of genotype, although severe KO groups start from a lower myelin area fraction stage.

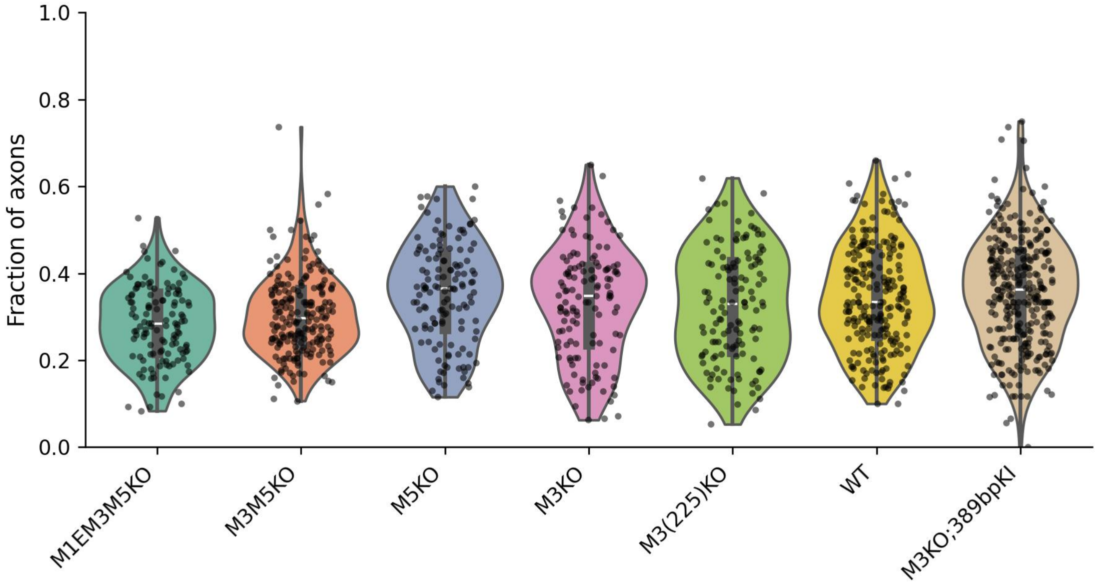


**Fig. S26. Fraction of axons with at least one mitochondria per-image for each *Mbp* genotype, pooled across ages and regions.**

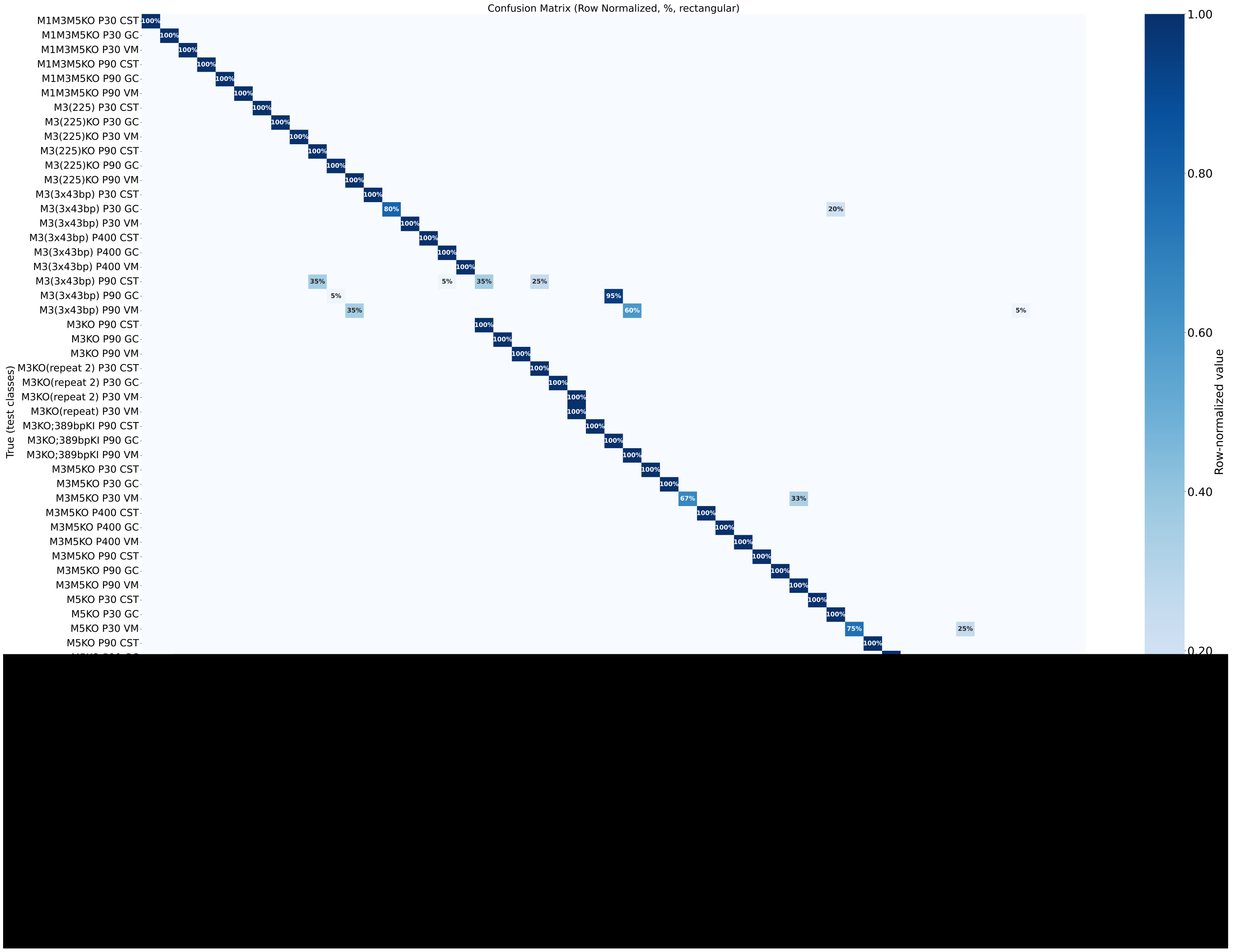


**Fig. S27. Cross-condition generalization evaluated using a leave-one-animal-out classification framework.** Rows represent held-out test conditions not included in training, and columns correspond to trained genotype classes. Values indicate row-normalized prediction probabilities. For readability, only informative cells are annotated: entries $\geq 5\%$; lower values are left unannotated. Due to the large number of classes displayed, the figure may appear visually dense at publication scale; readers are encouraged to zoom in to inspect finer details.

#### C. Supplementary Tables.

**Table 1. Per-image regression summary: mitochondrial vs myelin area fractions.**

| Gene | n | Slope | $R$ | $R^2$ | $p$-value |
|---|---|---|---|---|---|
| M1EM3M5KO | 118 | 1.226 | 0.292 | 0.085 | 0.00133 |
| M3M5KO | 194 | 2.548 | 0.326 | 0.106 | 3.56e-06 |
| M5KO | 124 | 2.458 | 0.301 | 0.091 | 0.000685 |
| M3KO | 120 | 3.880 | 0.556 | 0.309 | 4.48e-11 |
| M3(225)KO | 122 | 4.332 | 0.536 | 0.287 | 2.05e-10 |
| WT | 208 | 2.701 | 0.333 | 0.111 | 8.69e-07 |
| M3KOKI | 291 | 1.292 | 0.158 | 0.025 | 0.00688 |

**Table 2. Per-image regression summary: axon vs myelin area fractions.**

| Gene | n | Slope | $R$ | $R^2$ | $p$-value |
|---|---|---|---|---|---|
| M1EM3M5KO | 118 | 0.378 | 0.901 | 0.811 | 8.57e-44 |
| M3M5KO | 194 | 0.317 | 0.401 | 0.161 | 7.13e-09 |
| M5KO | 124 | 0.689 | 0.602 | 0.362 | 1.48e-13 |
| M3KO | 120 | 0.544 | 0.559 | 0.313 | 3.24e-11 |
| M3(225)KO | 122 | 0.408 | 0.500 | 0.250 | 4.51e-09 |
| WT | 208 | 0.439 | 0.380 | 0.145 | 1.46e-08 |
| M3KOKI | 291 | 0.284 | 0.298 | 0.089 | 2.13e-07 |

**Table 3. Regression summary for fiber diameter vs g-ratio (myelinated fibers only).**

| Age | Gene | Region | n | Slope | $R$ | $R^2$ | $p$-value |
|---|---|---|---|---|---|---|---|
| P30 | M1EM3M5KO | CST | 80 | -0.021 | -0.101 | 0.010 | 0.374 |
| P30 | M1EM3M5KO | GC | 258 | 0.040 | 0.269 | 0.073 | 1.15e-05 |
| P30 | M1EM3M5KO | VM | 184 | 0.095 | 0.474 | 0.225 | 1.03e-11 |
| P30 | M3KO | CST | 303 | 0.088 | 0.384 | 0.147 | 4.57e-12 |
| P30 | M3KO | GC | 405 | 0.088 | 0.421 | 0.177 | 8.09e-19 |
| P30 | M3KO | VM | 947 | 0.101 | 0.333 | 0.111 | 6.56e-26 |
| P30 | WT | CST | 350 | 0.055 | 0.197 | 0.039 | 2.03e-04 |
| P30 | WT | GC | 503 | 0.152 | 0.552 | 0.305 | 1.84e-41 |
| P30 | WT | VM | 941 | 0.080 | 0.281 | 0.079 | 1.56e-18 |
| P90 | M3KO | CST | 342 | 0.111 | 0.432 | 0.187 | 5.52e-17 |
| P90 | M3KO | GC | 294 | 0.123 | 0.500 | 0.250 | 4.95e-20 |
| P90 | M3KO | VM | 699 | 0.125 | 0.374 | 0.140 | 1.24e-24 |
| P90 | WT | CST | 553 | 0.077 | 0.325 | 0.106 | 4.23e-15 |
| P90 | WT | GC | 619 | 0.071 | 0.336 | 0.113 | 9.12e-18 |
| P90 | WT | VM | 849 | 0.111 | 0.407 | 0.166 | 3.61e-35 |
| P400 | WT | CST | 263 | 0.074 | 0.243 | 0.059 | 7.05e-05 |
| P400 | WT | GC | 311 | 0.245 | 0.667 | 0.445 | 1.96e-41 |
| P400 | WT | VM | 681 | 0.120 | 0.405 | 0.164 | 2.87e-28 |

**Table 4. Regression summary for fiber diameter vs mitochondrial area (myelinated fibers only).**

| Age | Gene | Region | n | Slope | $R$ | $R^2$ | $p$-value |
|---|---|---|---|---|---|---|---|
| P30 | M1EM3M5KO | CST | 42 | 0.089 | 0.491 | 0.241 | 0.002 |
| P30 | M1EM3M5KO | GC | 172 | 0.045 | 0.310 | 0.096 | 1.72e-04 |
| P30 | M1EM3M5KO | VM | 101 | 0.121 | 0.512 | 0.262 | 7.31e-08 |
| P30 | M3KO | CST | 87 | 0.040 | 0.326 | 0.106 | 0.0027 |
| P30 | M3KO | GC | 207 | 0.036 | 0.266 | 0.071 | 1.74e-04 |
| P30 | M3KO | VM | 431 | 0.061 | 0.475 | 0.226 | 2.11e-23 |
| P30 | WT | CST | 96 | 0.017 | 0.136 | 0.018 | 0.213 |
| P30 | WT | GC | 199 | 0.043 | 0.384 | 0.148 | 2.11e-08 |
| P30 | WT | VM | 464 | 0.066 | 0.540 | 0.292 | 2.08e-34 |
| P90 | M3KO | CST | 85 | -0.019 | -0.150 | 0.023 | 0.185 |
| P90 | M3KO | GC | 170 | 0.036 | 0.330 | 0.109 | 1.60e-05 |
| P90 | M3KO | VM | 341 | 0.115 | 0.592 | 0.350 | 1.59e-32 |
| P90 | WT | CST | 141 | 0.022 | 0.225 | 0.050 | 0.008 |
| P90 | WT | GC | 266 | 0.029 | 0.252 | 0.063 | 4.94e-05 |
| P90 | WT | VM | 404 | 0.042 | 0.232 | 0.054 | 3.32e-06 |
| P400 | WT | CST | 65 | -0.012 | -0.103 | 0.011 | 0.478 |
| P400 | WT | GC | 124 | 0.026 | 0.165 | 0.027 | 0.100 |
| P400 | WT | VM | 294 | 0.116 | 0.515 | 0.266 | 5.57e-19 |
